# Deletion of adipocyte Sine Oculis Homeobox Homolog 1 prevents lipolysis and attenuates skin fibrosis

**DOI:** 10.1101/2024.05.22.595271

**Authors:** Nancy Wareing, Tingting W Mills, Scott Collum, Minghua Wu, Lucy Revercomb, Rene Girard, Marka Lyons, Brian Skaug, Weizhen Bi, Meer A. Ali, Haniyeh Koochak, Anthony R Flores, Yuntao Yang, W Jim Zheng, William R Swindell, Shervin Assassi, Harry Karmouty-Quintana

**Author notes:** Co-corresponding authors Corresponding Authors: Shervin Assassi M.D., M.S. 6431 Fannin Street, Suite 5.260, Houston, TX 77030, Harry Karmouty-Quintana, Ph.D. 6431 Fannin Street, Suite 6.214, Houston, TX 77030.

## Abstract

Dermal fibrosis is a cardinal feature of systemic sclerosis (SSc) for which there are limited treatment strategies. This is in part due to our fragmented understanding of how dermal white adipose tissue (DWAT) contributes to skin fibrosis. We identified elevated sine oculis homeobox homolog 1 (SIX1) expression in SSc skin samples from the GENISOS and PRESS cohorts, the expression of which correlated with adipose-associated genes and molecular pathways. SIX1 localization studies identified increased signals in the DWAT area in SSc and in experimental models of skin fibrosis. Global and adipocyte specific Six1 deletion abrogated end-stage fibrotic gene expression and dermal adipocyte shrinkage induced by SQ bleomycin treatment. Further studies revealed a link between elevated SIX1 and increased expression of SERPINE1 and its protein PAI-1 which are known pro-fibrotic mediators. However, SIX1 deletion did not appear to affect cellular trans differentiation. Taken together these results point at SIX1 as a potential target for dermal fibrosis in SSc.

**Research in context:** *Evidence before this study:* Skin thickening and tightening are leading causes of morbidity in systemic sclerosis (SSc). The authors previously reported that the aberrantly expressed developmental transcription factor sine oculis homeobox homology 1 (SIX1) drives pulmonary fibrosis. However, the contribution of SIX1 to skin fibrosis and associated dermal fat loss remains unknown.

*Added value of this study:* The role of dermal fat loss in skin fibrosis is not fully understood. Studies have shown that adipocytes can transition to mesenchymal cells promoting fibrosis, consistent with loss of the dermal white adipose layer. Our research provides insight into a novel molecular mechanism of lipodystrophy important for skin fibrosis in SSc. We identified the upregulation of *SIX1* in adipocytes in skin from patients with SSc which was associated with the progression of skin fibrosis. We found elevated *Six1* in mouse dermal adipocytes of early fibrotic skin. Ubiquitous and adipose-specific loss of *Six1* decreased markers of experimental skin fibrosis in mice which recapitulate cardinal features of SSc dermal fibrosis. Increased SIX1 expression is linked with elevated levels of Serpine1 the gene that codes for the protein plasminogen activator inhibitor (PAI)-1. This is important since PAI-1 is a known pro-fibrotic agent in the skin that contributes to the deposition of extracellular matrix (ECM) products.

*Implications of all the available evidence:* Fat atrophy may represent a targetable contributor to early systemic sclerosis manifestations. This is as it precedes skin fibrosis and the use of topical agent which are usually lipophilic can help us target dermal adipocytes. Our results show that SIX1 could be an important early marker for skin fibrosis in SSc that can also be targeted therapeutically.

## Introduction

Systemic sclerosis (SSc; or scleroderma) is a rare and heterogenous autoimmune connective tissue disorder.^1,2^ The multiorgan dysfunction has been characterized as a triad of immune dysregulation, vasculopathy, and excessive extracellular matrix (ECM) deposition by myofibroblasts, leading to skin and internal organ fibrosis.^2–4^ Skin thickening and tightening is responsible for considerable morbidity in this debilitating disease.^5,6^ The extent of skin fibrosis defines the two subclasses of SSc: limited cutaneous (lcSSc) and diffuse cutaneous SSc (dcSSc).^3^ There are currently no FDA-approved treatments for skin involvement in SSc. More extensive skin involvement at the time of diagnosis is associated with higher levels of disability, severity of pain, and decreased survival.^7,8^ This underscores the urgent need to identify novel targeted therapies for the treatment of SSc.^9^

Thus, understanding the early pathogenesis of disease represents a critical step towards novel therapeutic approaches.^9^ The mechanisms that lead to organ fibrosis in SSc are not fully understood, however, similar mediators have been identified to play a role in the lung and skin. For example, increased adenosine, hyaluronan and IL-6 have been implicated in the pathophysiology of both lung and skin fibrosis^10–14^. However, these mechanisms have largely focused on fibroblast and epithelial biology, with limited studies in adipocytes. This is important as an early hallmark of SSc is skin-associated adipose tissue atrophy and replacement by extracellular matrix leading to dermal thickening.^15,16^ Clinically, this may directly contribute to rigidity and tethering observed in early lesional SSc skin.^15,16^. Dermal white adipocyte tissue (DWAT) adipocytes display highly distinct features compared to other white adipocytes, including significant plasticity.^17–19^ They can cycle through de-differentiation and re-differentiation as part of a physiologic response to hair cycling, aging, and energy demands.^17,18,20^ Under certain conditions, de-differentiated pre-adipocytes escape the normal cycle of redifferentiation, and *trans*-differentiate into ECM-producing myofibroblasts. This process is referred to as the adipocyte-to-myofibroblast transition (AMT).^21,22^ AMT has been well-studied in wound healing ^21,23^ and recent evidence suggests it may also contribute to pathological skin fibrosis.^22,24^ Mature adipocytes in the skin produce PDGF ligands and BMPs, both of which are implicated in wound healing and fibrosis. ^25^ In addition the adipose secretome has also been identified to play a role in the pathophysiology of SSc. ^26^ Lipid-filled adipocytes cross-talk with other cell types in the stromo-vascular fraction of adipose tissue. The interaction between the dermal fibroblasts and adipocytes has also been appreciated as a contributor to irregular inflammation and aberrant wound healing through pro-inflammatory signaling between cell types ^27^ and adipocyte-driven regulation of fibroblast ECM production. ^28^ A robust physiologic axis also exists between adipocytes and endothelial cells, by which cell signaling can be bi-directionally regulated. ^29^ However, mechanisms by which adipocytes contribute to the pathogenesis of skin fibrosis in SSc remain poorly understood.

Our group and others identified sine oculis homeobox homolog 1 (SIX1) as a novel mediator in lung fibrosis promoting release of pro-fibrotic mediators by alveolar epithelial cells.^30–32^ Further, SIX1 has also been implicated in asthmatic lung fibrosis ^31,32^ and in liver fibrosis defined by excessive myofibroblast activation and ECM deposition ^33^. Lung fibrosis is an important complication of SSc that is typically observed following onset of skin fibrosis and while epithelial and fibroblast based mechanisms are most highly studied, the contribution of the adipocyte to fibrotic process is not fully known.^34^ However, whether SIX1 plays a role in skin fibrosis in SSc is not known, in particular how adipocyte SIX1 regulates dermal fibrosis remains under investigated. SIX1 is a member of an evolutionarily conserved family of developmental transcription factors.^35^ SIX1 plays a critical role in regulating expression of genes that control precursor cell survival and proliferation during embryogenesis. In healthy adults, SIX1 is negligibly expressed in most tissues.^36^ Perhaps the most well-studied role of SIX1 in adulthood is in the context of cancer where it is a critical regulator of trans-differentiation of pre-cancerous cells into mesenchymal cells with metastatic features.^37–41^ Although *SIX1* transcript expression has been identified in healthy subcutaneous adipose tissue^42^, to date it’s exact role in adipose tissue biology remains poorly investigated. Brunmeir et al.^43^ were the first to identify the direct transcription regulation by SIX1, and interaction between SIX1 and major regulators of adipogenesis, in mature fat cells. Recently, it was demonstrated that *in vivo* SIX1 overexpression in mouse hepatocytes exacerbate diet-induced liver inflammation, metabolic disruption, and hepatic steatosis, and activates liver-specific receptors to induce de novo lipogenesis.^44^ This work is founded upon the fundamental and newly developing understanding of the roles of SIX1, particularly in the realm of lipolysis and the release of pro-fibrotic factors that regulate dermal fibrosis. We hypothesize that SIX1 contributes to dermal lipoatrophy and skin fibrosis in SSc. Surprisingly, our data suggests that SIX1 is not involved in modulating adipocyte phenotype but instead it regulates the release of the pro-fibrotic mediator, PAI-1 that promotes dermal fibrosis.

## Methods

### Study populations

#### GENISOS cohort

The prospective cohort study, GENISOS (Genetics versus ENvironment In Scleroderma Outcome Study), is a collaboration between The University of Texas Health Science Center Houston, The University of Texas Medical Branch at Galveston and the University of Texas Health Science Center at San Antonio. All participants meet the diagnosis of SSc according to the American College of Rheumatology preliminary classification.^45^ Details of recruitment and selection criteria have been previously published.^46^ Individuals with diffuse cutaneous (dc) SSc and limited cutaneous (lc) SSc are enrolled within five years of disease onset, defined as the first non-Raynaud’s symptom. The study was approved by the institutional review board of all participating sites, and written informed consent was obtained from all study subjects.

The modified Rodnan skin score (mRSS) was calculated by a board-certified rheumatologist with extensive experience in assessment of scleroderma skin ^47^. The mRSS is determined by assessment of the skin thickness of 17 body areas by physical examination, The mRSS serves as a surrogate for disease activity, severity and mortality in patients with SSc ^48^. Healthy control individuals are enrolled to serve as controls. SSc-affected individuals and controls are matched at a ratio of 3:1 based on age, sex, and ethnicity. Gene expression analysis from SSc-affected skin and skin from controls has been previously described.^49^ Briefly, global gene expression is assessed using Illumina HumanHT-12 bead array. Raw data were analyzed with BRB ArrayTools. 113 SSc-affected individuals and 44 unaffected controls had available *SIX1* expression levels.

#### PRESS cohort

The Prospective Registry for Early Systemic Sclerosis (PRESS) cohort is a multi-site observational cohort of individuals with dcSSc enrolled within three years of onset of first non-Raynaud’s symptom.^9^ All participants fulfill the 2013 American College of Rheumatology (ACR)/European League Against Rheumatism (EULAR) classification criteria for SSc.^50^ RNA sequencing data from the skin of PRESS participants and controls, previously utilized by our group,^51^ was queried for expression of *SIX1*. Forty-eight SSc-affected individuals and thirty-three controls had available *SIX1* expression levels. The R Bioconductor package edgeR6 analysis was utilized to identify differentially expressed transcripts between SSc patients and healthy controls with a false discovery rate cutoff of 0.05 and fold change cutoff of >1.5 or <0.67

### Cell type-specific expression signatures

Cell type-specific expression signatures were originally developed as previously described ^52^ and have been utilized by our group.^51^ A “cell-type specific signature score” denotes a set of genes for which expression in a given cell type is notably higher than expression in the other evaluated cell types. The numerical value of each score was calculated based on fold-change estimates (SSc versus control) for 125 “signature genes” of a given cell type. The methodology used is described in detail in ^49,52,53^, briefly for gene expression analysis data were imported into BRB-ArrayTools as processed signal values. Values were excluded if the mean signal was not significantly greater than background. Values were then log2 transformed followed by quantile normalization. Genes with >20% missing values across arrays were filtered out. The remaining gene values were used for analyses.

### Pathway Analysis

Herein we used the same protocol described used previously^49,52,53^, genes that were differentially expressed on average in SSc compared to control at a false discovery rate (FDR) of <0.05 were uploaded to Ingenuity Pathway Analysis (Qiagen). Expression analysis was performed using “Human Genome CGH 44K” as reference set, including “direct and indirect relationships,” with Confidence set to “Experimentally observed” only. The analysis was then repeated with genes that were differentially expressed on average in SSc compared to control at an FDR of <0.1.

### Correlation analyses

The correlation between signature score and *SIX1* expression was evaluated for each sample evaluated, then the mean Spearman’s rank correlation co-efficient was reported. Individual gene correlations were analyzed by Spearman’s rank correlation. Correlation coefficients are reported as “r.”

### Functional annotation of all differentially expressed genes in PRESS

All differentially expressed genes (DEGs) in the skin biopsies of PRESS cohort participants with SSc were input into The **D**atabase for **A**nnotation, **V**isualization and **I**ntegrated **D**iscovery (DAVID) for functional annotation (Available at https://david.ncifcrf.gov/summary.jsp). A Bonferroni cut-off of 0.5 was used to determine significantly enriched biological pathways.

### Animal studies

All studies were reviewed and approved by UTHealth Houston Animal Welfare Committee (AWC-19-0029 and AWC-22-0028). Six to eight-week-old male and female C57/BL6J mice were used for experiments with wild-type mice. Mice with dorsal skin in the telogen phase of hair cycle were used. Regions of skin in the anagen phase were excluded. When possible, littermates were equally distributed between groups. Group sizes were determined by power analysis with an alpha of 0.05 and a power of 80%.

#### Subcutaneous bleomycin model

The shaved dorsum of isoflurane-anesthetized mice was injected subcutaneously with phosphate-buffered saline (PBS; vehicle) or 0.1 Units/ml bleomycin sulfate USE ([bleo]; Teva Pharmaceuticals) diluted in PBS. For up to 28 days, 0.02 Units of bleo per mouse per day was distributed between two sites upper and lower dorsal (100µl per site) 6 days per week as previously described ^54^. The upper dorsal section was utilized for RT-qPCR and the lower dorsal for histological analyses. The day following the last injection, mice were euthanized with CO_2_ followed by cervical dislocation. Any mice in the anagen stage of the hair cycle were excluded from final analyses.

#### Transgenic mouse models

Tamoxifen-inducible ubiquitous *Six1* knock-out mice (iUbc-Six1^-/-^) were generated by cross-breeding B6.Cg-Ndor1^Tg(UBC-cre/ERT2)1Ejb^/1J (iUbc^Cre^) (Jackson Labs Cat No. 007001; a gift from lab of Holger Eltzschig, McGovern Medical School, Houston, Texas, USA) and Six1^loxP/loxP^ (a gift from Pascal Maire, Inserm U1016, Institut Cochin. 75014 Paris, France). Tamoxifen-inducible adipocyte-specific *Six1* knock-out mice (iAdipo-Six1^-/-^) were generated by cross breeding C57BL/6-^Tg(Adipoq-cre/ERT2)1Soff/J^ (iAdipo^Cre^) (Jackson Labs; Cat No. 025124) and Six1^loxP/loxP^ mice. Six1^loxP/loxP^ mice contain loxp sites within the 5’ untranslated region of exon 1 and the intronic sequence between exon 1 and exon 2 of the *Six1* gene.^55^ iUbc^Cre^ and iAdipoCre mice express a Cre recombinase-mutated estrogen receptor fusion protein (Cre-ERT2) under transcriptional control of the human *Ubiquitin C/Ubc* promoter or the mouse *Adiponectin/Adipoq* promoter, respectively.^56^ When bound by tamoxifen, Cre-ERT2 translocates from the cytoplasm to the nucleus, targets loxp sites, and efficiently deletes the floxed allele.^57^ All mouse strains have a C57BL/6 background.

#### Tamoxifen treatment

Mice were administered tamoxifen (Sigma-Aldrich, Cat# T5648) dissolved in corn oil (20mg/ml) shaken overnight at 37°C. Mice were intraperitoneally administered 75mg/kg tamoxifen in corn oil every 24 hours for 5 days. Mice were then monitored for 7 days prior to start of the experiment (Protocol from The Jackson Laboratory; https://www.jax.org/research-and-faculty/resources/cre-repository/tamoxifen).

#### Tissue processing

Dorsum of mice were shaved, depilated with Nair, then cleaned and allowed to air dry. Two 8mm punch biopsies were taken from sites of injection. The inferior biopsy was immediately placed in 10% neutral buffered formalin and stored at 4°C for at least 48 hours and no longer than 1 week. Formalin-fixed tissue was embedded in paraffin. The superior biopsy was cut in half. One half was placed in cold RNAlater (Sigma-Aldrich, Cat# R0901) and stored per manufacturer’s instructions. For RNA isolation, frozen tissue was finely minced and placed in QIAzol Lysis Reagent (Qiagen, Cat# 79306). Tissue was homogenized on ice using the TissueRuptor (Qiagen, Cat# 9002755) and homogenate was incubated at room temperature for 5 minutes. RNA extraction was performed using the miRNeasy Mini Kit (Qiagen, Cat# 217004) according to manufacturer’s instructions. RNA was eluted and stored at -80°C until use.

### Reverse transcription quantitative polymerase chain reaction (RT-qPCR)

RNA quality was measured by bioanalyzer using RNA 600 Nano Assay (Agilent, Part No 5067-1511). RNA with an RNA Integrity Number > 7 was used for downstream experiments. RNA concentration was measured by nanodrop. Reverse transcription was performed using the QuantiTect Reverse Transcription Kit (Qiagen, Cat#205313) with genomic DNA removal. PCR was carried out with iTaq Universal SYBR Green Supermix (Bio-Rad, Cat#1725124). A complete list of SYBR green primers is listed in **<u>Supplementary</u> Table 1**. qPCR was performed using BioRad CFX Opus 384 Real-Time PCR System. Data processing was performed on BioRad CFX Maestro. Expression values are reported as 2^-ΔCt^ (Ct_gene of interest_ – Ct_18s rRNA_).

### Immunofluorescence

Five-micron thick formalin fixed paraffin-embedded (FFPE) skin tissue was deparaffinized then heat-based antigen retrieval was performed in high pH Tris-based antigen retrieval buffer (Vector Laboratories, Cat# H-3301). After washing in tris-buffered saline containing 0.1% Tween-20 (TBS-T), tissue was blocked in 2.5% - 10% animal serum for 1 hour at room temperature. Primary antibodies were perilipin 1 (Abcam #ab3526, 1:200), collagen 6 (Abcam #ab182744, 1:100). Primary antibodies were incubated overnight at 4°C. After washing in TBS-T, tissue was incubated with Alexa Fluor secondary antibody (ThermoFisher Scientific) for 1 hour at room temperature. Autofluorescence was quenched using TrueVIEW Autofluorescence Quenching Kit (Vector Laboratories, Cat# SP-8400-15). Tissue was mounted using ProLong Gold Antifade Mountant with DAPI (Invitrogen, Cat# P36931).

### *In situ* hybridization

RNAScope, an optimized single-molecular *in situ* hybridization technique ^58^ was used to detect single RNA transcripts in FFPE human and mouse skin tissue. RNAScope 2.5 HD Assay – RED (Advanced Cell Diagnostics; Cat No. 322360) and RNAScope 2.5 HD Duplex Reagent Kit (Advanced Cell Diagnostics; Cat No. 322430) were used for detection of one or two transcripts, respectively. Complete protocol is available from manufacturer. RNAScope probes are listed in <u>**Supplementary Table 2**</u>.

### Imaging and histological analysis

Histology images were captured on the Keyence BZ-X810 microscope and analyzed using Keyence BZ analyzer software. Adipocyte droplet size was measured by calculating the area within the droplet as outlined by cytoplasmic perilipin 1 staining. Collagen 6 immunofluorescence was quantified as percent positive area of the DWAT or brightness intensity within the DWAT. A macro was created using the Hybrid Cell Count feature to ensure unbiased and consistent quantification parameters. DWAT was defined by the region containing positive perilipin 1 staining and the upper border of the panniculus carnosus. Hair follicles and large vessels were excluded. Four to five non-continuous images per mouse biopsy were used for all histological analyses.

### In Vitro Cell Culture Studies

Low-passage 3T3-L1 cells (ATCC Cat#CL-173) were thawed and to 70% confluency in 10ml of complete pre-adipocyte growth media composed of high glucose Dulbecco’s Modified Eagle’s Medium (DMEM) containing 4mM L-glutamine (Sigma Aldrich, Cat# D5796) supplemented with 10% calf serum (Colorado Serum Company, Cat#31334), 1% penicillin and streptomycin (Gibco, Cat# 15140-148), and 1mM sodium pyruvate (Sigma Aldrich, Cat# S8636). Cells were washed in PBS and media was replaced every 72 hours. To prevent growth arrest, expanding pre-adipocytes were never allowed to reach greater than 70% confluency. Cells were detached using 0.25% trypsin-EDTA.

To induce pre-adipocyte differentiation into mature white adipocytes, previously described protocols were utilized^59,60^ and the cells were allowed to grow until day 9. Control siRNA (Sigma SIC001) or small interfering siRNA for Six1 (Sigma SASI_Mm01_00198105) was transfected to 3T3 cells on days -4, -2 of differentiation to mature fibroblasts. For RNA isolation, 85µl per cm2 of cold Trizol (Invitrogen, Cat# 15596026) was used. For protein isolation, 8µl per cm^2^ of cold RIPA buffer with protease and phosphatase inhibitors (ProteinBiology, Cat# 78442) was used. For immunoblots, cells were lysed with RIPA buffer(Thermo Scientific) containing Halt Protease and Phosphatase inhibitor(Sigma Aldrich). Samples were loaded in reducing Sodium dodecyl-sulfate (SDS) buffer (BD Biosiences) in 4-20% Tris-Glycine eXtended (TGX) gels (Bio-Rad). Polyvinylidene difluoride (PVDF) membranes were blocked with 5% dry milk in TBST for 1 hr and then incubated overnight with primary antibody. HRP conjugated secondary antibody was incubated for 1 hour and then blots were imaged with SuperSignal West Pico (Thermo Scientific) on a Chemi-Doc Imaging system(Bio-Rad). Antibodies used included: β-Actin(Cell Signaling Technology [CST] 5125S at 1:2000); SERPINE1 (abcam, ab282007 at 1:1000); SIX1 (CST 12891S at 1:1000) and Rabbit HRP (CST 7074S 1:3000).

### NanoString nCounter Analysis System

After RNA extraction from samples, their quality control was done using Agilent TapeStation, and the amount of RNA was quantified with Invitrogen™ Qubit™ RNA High Sensitivity (HS) kit. Later, the samples went through an overnight hybridization process --performed at 65°C in a thermocycler--where RNA (analytes) were hybridized to nCounter optical barcodes (nCounter® Mouse Fibrosis V2 Panel: NanoString Technologies). After hybridization, samples were processed using the nCounter Pro Analysis System which rapidly immobilizes and counts samples that have hybridized to nCounter barcodes. This is an ex-situ digital counting of RNAs. Briefly, on the nCounter Prep Station, hybridized samples get prepared for data collection on the nCounter Digital Analyzer. This preparation includes purification and immobilization in a sample cartridge. On the nCounter Digital Analyzer, hundreds of images are collected for each sample, yielding hundreds of thousands of target molecule counts per sample. After an internal processing of images, the results are RCC (Reporter Code Count) files containing the RNA counts that can be directly processed through the nSolver Analysis Software.

### Statistical analysis

Prism software (v9.0; GraphPad or higher) was used for all statistical analyses. ROUT outlier test was performed on all datasets. Outliers were excluded if FDR greater than 1%. Two-tailed t-test with Welch’s correction was used for 2 group comparisons. Two-way ANOVA with multiple comparisons and correction using the Sidak method was used for 3 or more groups. Detailed statistical analysis for each experiment is shown in the figure legends.

## Results

### Increased SIX1 levels correlate to skin fibrosis in SSc patients

Previously our group identified increased SIX1 in lung fibrosis^30^ and SIX1 expression has been identified to be present in subcutaneous adipocytes particularly in those exhibiting aberrant function ^42^. As such we aimed to determine whether SIX1 was elevated in skin samples from SSc patients. To do this, we first determined expression of SIX1 from two distinct cohorts: The GENISOS cohort that includes patients with lcSSc and dcSSc at different stages of disease and the PRESS cohort, enriched for patients with early-stage dc-SSc. *SIX1* transcript levels were elevated in SSc skin in limited SSc, diffuse SSc, and in early diffuse SSc compared to control skin in both independent cohorts (<u>**Figure 1**</u>). In the PRESS cohort, that is enriched for patients with early dcSSc, RNA-sequencing revealed increased SIX1 signals (**Figure 1a**). In the GENISOS cohort, which encompasses dcSSc and lcSSc at different stages of disease, *SIX1* signal intensity, denoting expression levels was higher in patients with dcSSc compared to lcSSc (**Figure 1b**). To determine whether *SIX1* is associated with the genomic landscape of a particular cell type, we correlated *SIX1* expression levels with cell type-specific signature scores previously utilized by our group.^51,61^ The bioinformatic analysis identified genes that are expressed at comparatively higher levels in a specific cell type and created a signature score for each cell type being evaluated. The subcutaneous adipose signature was the most highly correlated with *SIX1* expression in both the GENISOS (r = 0.76) and PRESS cohorts (r = 0.79) this point at SIX1 as an important mediator that is elevated in adipose tissue in SSc (**Figure 1c**). Higher expression of genes specific to fibroblasts, vascular and lymphatic endothelial cells significantly correlated with higher expression of *SIX1* suggesting that SIX1 may regulate mesodermal derived cells during fibrosis but not the ectodermal derived epithelium. Individual gene correlation analysis revealed that genes associated with adipocyte biology were enriched amongst those genes most highly correlated with *SIX1* specifically in early dcSSc skin (**Figure 1d; <u>Supplementary Table 3</u>**). These included genes encoding proteins required for adipocyte differentiation and triglyceride metabolism, including *ADIPOQ* and *PPARG*.^62–64^ Functional annotation of all DEGs in the skin of dcSSc patients in the PRESS cohort compared to healthy controls showed significant enrichment for “regulation of lipolysis in adipocytes” in addition to “ribosome” and “AMPK signaling pathway” (**Figure 1e**).

**Figure 1.**
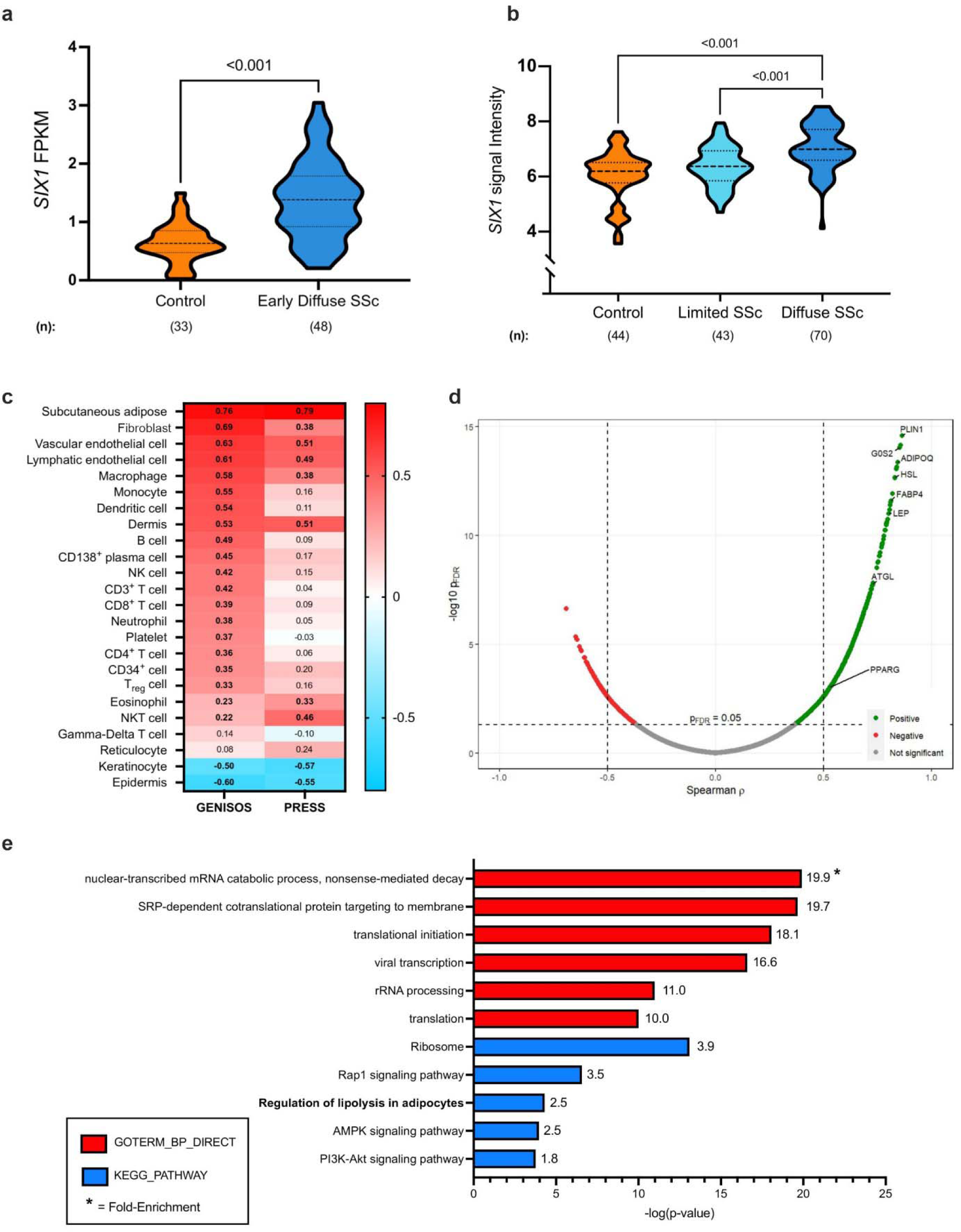
*SIX1* is elevated in SSc skin and correlates with adipose-related genes and pathways. **a**) *SIX1* expression in PRESS SSc skin samples and controls. FPKM = fragments per kilobase million based on a false discovery rate cutoff of 0.05 and fold change cutoff of >1.5 or <0.6 **b**) *SIX1* expression in baseline SSc skin samples and controls in the GENISOS cohort based on Student’s t-test **c**) Heatmap showing correlation (“r”) between skin *SIX1* expression and cell type signature scores on Spearman’s Rank order correlation. Bolded pathways indicate correlations with p-values <0.05 in both cohorts. **d**) Volcano plot showing individual gene-*SIX1* correlations in PRESS cohort SSc skin samples. PLIN1=perilipin 1; G0S2=G0/G2 switch gene 2; ADIPOQ=adiponectin; HSL=hormone sensitive lipase; FABP4=fatty acid binding protein 4; LEP=leptin; ATGL=adipose triglyceride lipase; PPARG= peroxisome proliferator activated receptor gamma based on a Spearman’s rank coefficient analysis. **e)** KEGG pathway annotation of all DEGs in SSc skin of PRESS cohort participants using GoStats. Significance levels *** P ≤ 0.001 refer to a Bonferroni cut-off of 0.5 analysis (panel a) or a the R Bioconductor package edgeR6 analysis for to identify differentially expressed transcripts between SSc patients and healthy controls with a false discovery rate cutoff of 0.05 and fold change cutoff of >1.5 or <0.67 for panel b.

### Increased adipocyte SIX1 levels correlate with loss of dermal white adipose tissue (DWAT) in SSc

Several studies have shown that the lipoatrophy and loss of DWAT is present in skin fibrosis in SSc^15^ and even precedes the fibrotic matrix deposition^65^. In addition, AMT ^22,24^ and the adipose secretome ^26^ has been implicated in the pathophysiology of SSc. Thus, we next aimed to determine whether SIX1 levels were increased in the diminishing DWAT layer in the skin. Herein we assessed tissue samples from the GENISOS cohort that include SSc-affected skin samples with varying disease durations were compared to an age, sex, and ethnicity matched control samples which demonstrate a progressive loss of DWAT areas as disease progresses (**Figure 2a**). This is important as DWAT levels are known to reduce as dermal fibrosis develops, thus the GENISOS cohort allows us to temporally assess loss of adipose tissue and SIX1 levels during progression of disease. Demographic and clinical features of these individuals are listed in <u>**Supplementary Table 4**</u>. Compared to the control biopsy, there was notably less DWAT in the skin of all dcSSc-affected individuals, with the patient with the most established form of disease presenting with no discernible DWAT areas (**Figure 2a**). Thus, having established a correlation between changes in adipocyte biology, stromal cells and *SIX1* gene expression in SSc skin; we sought out to identify those cell types which produce *SIX1* in SSc skin with abnormal dermal fat.

**Figure 2.**
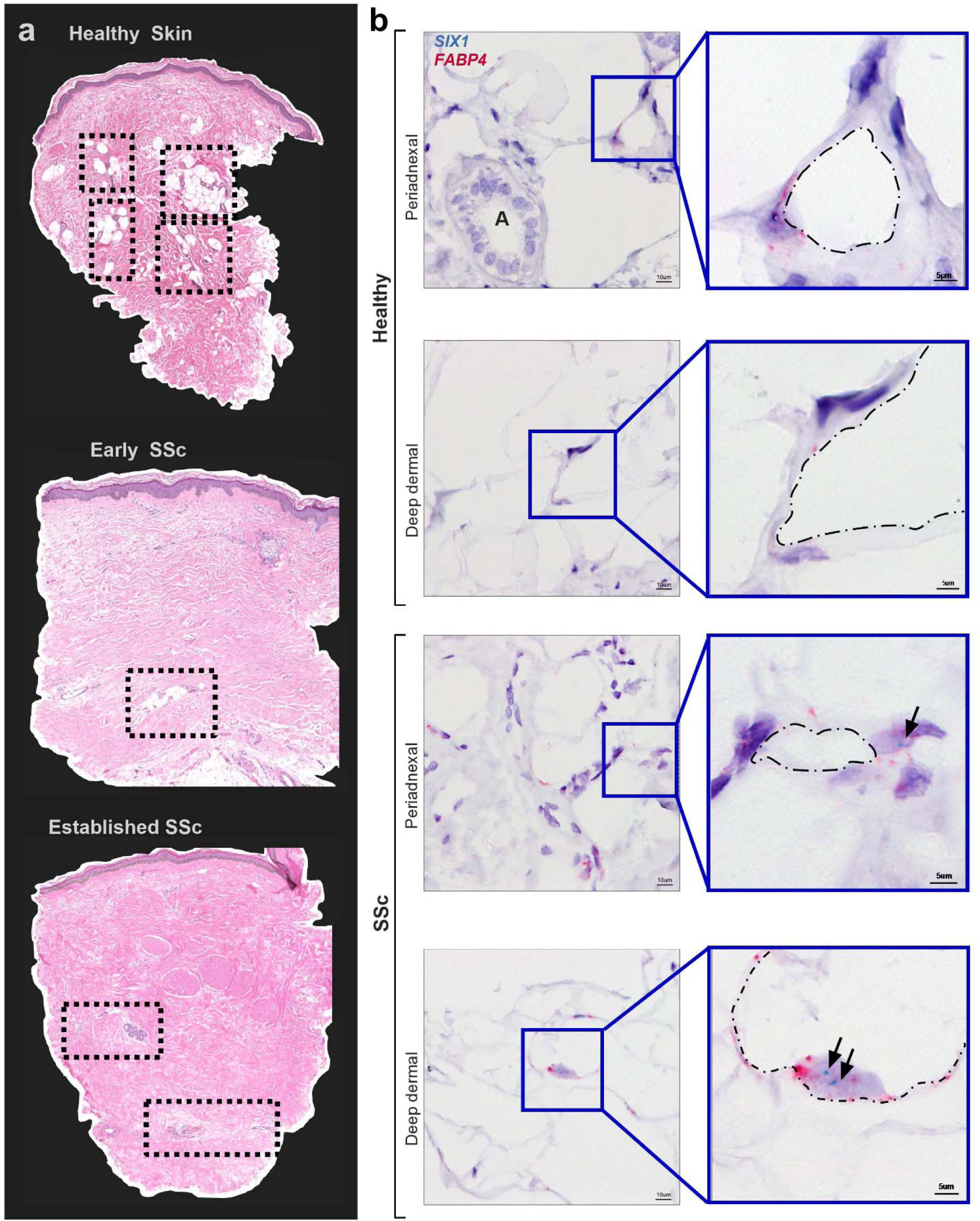
Dermal White Adipose Tissue (DWAT) atrophy and increased adipocyte SIX1 expression in SSc. **a)** Hematoxylin and eosin staining of human skin biopsies from GENISOS cohort participants. **top** Control skin. **middle** early SSc-representative image **bottom** established representative SSc-skin sample. Clinical and demographic features of biopsies individuals are provided in Supplementary Table 4. Boxes contain dermal white adipose tissue. Dotted boxes denote DWAT areas. **b)** Representative images of dual *in situ* hybridization for *SIX1* (teal) and *FABP4* (pink) in 3 SSc and 1 demographically matched control biopsies. Hematoxylin (purple) co-stain labels nuclei. “A” marks a dermal adnexal structure. Black arrows point to *SIX1* transcript signal. Dash dot outlines denote the interior periphery of adipocytes.

We selected 15 systemic sclerosis (SSc) patients who retained dermal fat, most of whom were within three years of developing diffuse disease. Additionally, skin biopsies from 5 healthy controls were included. To localize the SIX1 gene in these samples, we employed single molecule in situ hybridization. DWAT in human skin is localized around adnexal glands and hair follicles, and within and below the deep dermis^66^. *SIX1* was detected in both peri-adnexal and deep dermal adipocytes. Mature adipocytes are identifiable by a single-large lipid droplet surrounded by a thin ring of cytoplasm and a peripheral nucleus which expresses Fatty Acid Binding Protein 4 (*FABP4)* (<u>**Figure 2b**</u>). We acknowledge that some fibroblasts also express *FABP4*, however the unique morphology of adipocytes as described above make them easily distinguishable from the small, spindle-shaped fibroblast with a dominant, central nucleus. The clinical relevance of these findings was supported by positive correlations between the expression of *SIX1* in SSc skin, and the extent and severity of SSc skin fibrosis. Spearman’s rank-order correlation analysis showed positive correlation between whole skin *SIX1* expression and mRSS (r=0.40, p<0.001), and local skin score (r=0.38, p<0.001) near the site of the biopsy. To our knowledge, dermal fat *SIX1* has not previously been identified *in situ*, the significance of which is supported by clinical data linking *SIX1* to more severe and extensive SSc skin involvement. Together, these data from two SSc study cohorts provided a strong premise for investigating *SIX1* in SSc disease mechanisms.

### Adipose tissue loss is evident in a bleomycin model of dermal fibrosis

We selected the murine subcutaneous (SQ) bleo model of skin fibrosis^54^ as the pre-clinical model to study the effects of *SIX1* on fibrosis and lipoatrophy.^11,67^ Unlike human skin, rodent skin DWAT is separated from the SWAT by a thin layer of skeletal muscle (the panniculus carnosus), allowing us to distinguish dermal adipocytes from subcutaneous adipocytes without the use of additional markers.^68,69^ Further the fibrotic changes that occur over the 28 days of SQ bleo treatment allows us to study the role and expression of SIX1 during the pathogenesis of dermal fibrosis.^22,65,70^

We demonstrated that, serial injections of SQ bleo recapitulated lipodystrophy and dermal sclerosis mirroring SSc manifestations (**<u>Figure 3a</u>)**.^22,64,69^ Using Masson’s trichrome staining, we showed progressive bleo-induced atrophy of the DWAT, increased collagen deposition, and dermal thickening observable at weekly time points up to 28 days (**Figure 3b**). Attrition of DWAT was appreciable on histology as early as day 7 of bleo treatment; when dermal thickening was less pronounced. Quantification of dermal thickness and DWAT area confirmed these observations. When compared to day 7, dermal thickening was significant after 28 days of bleo (**Figure 3c**). This change lagged behind the significant decline in DWAT area (**Figure 3d**). Transcriptomic analysis revealed increased signals for *Six1* (**Figure 3e**) and despite minimal histological changes in the dermis after 7 days of bleo, prominent extracellular matrix genes collagen 1a1 (Col1a1), collagen 1a2 (Col1a2), and collagen 6a1 (Col6a1), transforming growth factor beta (*Tgfb1*) and of serine proteinase inhibitor E 1 (*Serpine1*) were upregulated (**Figure 3f-j**). Intriguingly we did not detect increased expression for peroxisome proliferator-activated receptor gamma (*Pparg) on day 7 of SQ bleo (***Figure 3k**). These results support that a reduction in DWAT and upregulation of fibrotic genes precedes significant dermal expansion following SQ bleo treatment. Next, we investigated whether adipocyte *Six1* expression, as observed in SSc skin, was recapitulated in the SQ bleo model. Dual in situ hybridization probed for skin *Six1* and *Adipoq* transcript expression after 7 days of SQ vehicle or bleo injections. *Adipoq* is highly and specifically expressed in lipid-laden adipocytes.^63^ Mirroring findings in human SSc skin whereby *SIX1* was elevated in early disease, *Six1* transcript was detectable in the dermal adipocytes after 7 days of SQ bleo treatment (<u>**Figure 4**</u>). *Six1* was minimally expressed in DWAT adipocytes in vehicle-treated skin (**Supplementary Figure 1**). Collectively, this data demonstrates that adipocyte loss is evident prior to fibrotic deposition and that SIX1 levels are increased in adipocytes by day 7 of bleo treatment.

**Figure 3.**
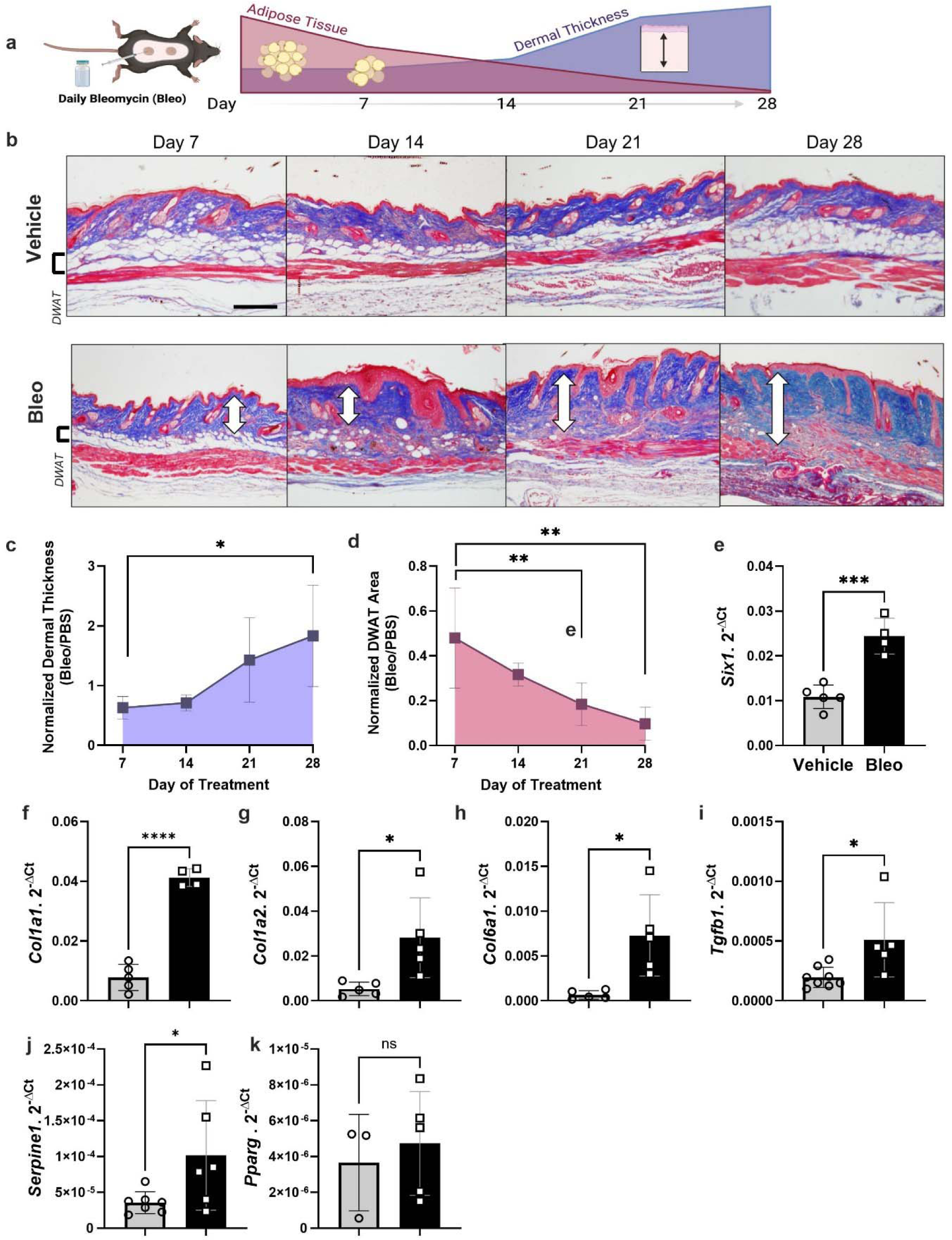
Dermal white adipose loss precedes fibrosis in the murine model of bleomycin induced skin fibrosis. **a**) Schematic representation of SQ bleo model of skin fibrosis in mice (Created with BioRender) **b)** Representative images of Masson’s trichrome staining of dorsal mouse skin after 7, 14, 21, and 28 days of SQ vehicle or bleo treatment. White arrows indicate dermal thickness. Scale bar = 200µm. **c)** Dermal thickness and **d)** area of DWAT at 7, 14, 21, and 28 days reported as the mean ±SD of bleo-injected mice normalized to the average of all vehicle-injected mice at that time point N=5 for each time point. Transcript expression levels for sine oculis homeobox homolog 1 (*Six1,* **e**),collagen 1a1 (*Col1a1,* **f**), collagen 1a2 (*Col1a2,* **g**), collagen 6a1 (*Col6a1,* **h**), transforming growth factor beta 1 (*Tgfb1,* **i**), serpin family E member 1 (*Serpine1*, **j**) and peroxisome proliferator activated receptor gamma (*Pparg,* **k**) in 7-day skin samples by RT-qPCR. Expression was normalized to 18s rRNA. DWAT = dermal white adipose tissue. Significance levels * P ≤0.05, ** P ≤0.01, *** P ≤0.001, and **** P ≤0.0001 refer to a simple linear regression for panels c-d and an unpaired t-test for panels e-k. Each individual plot represents a biological N number.

**Figure 4.**
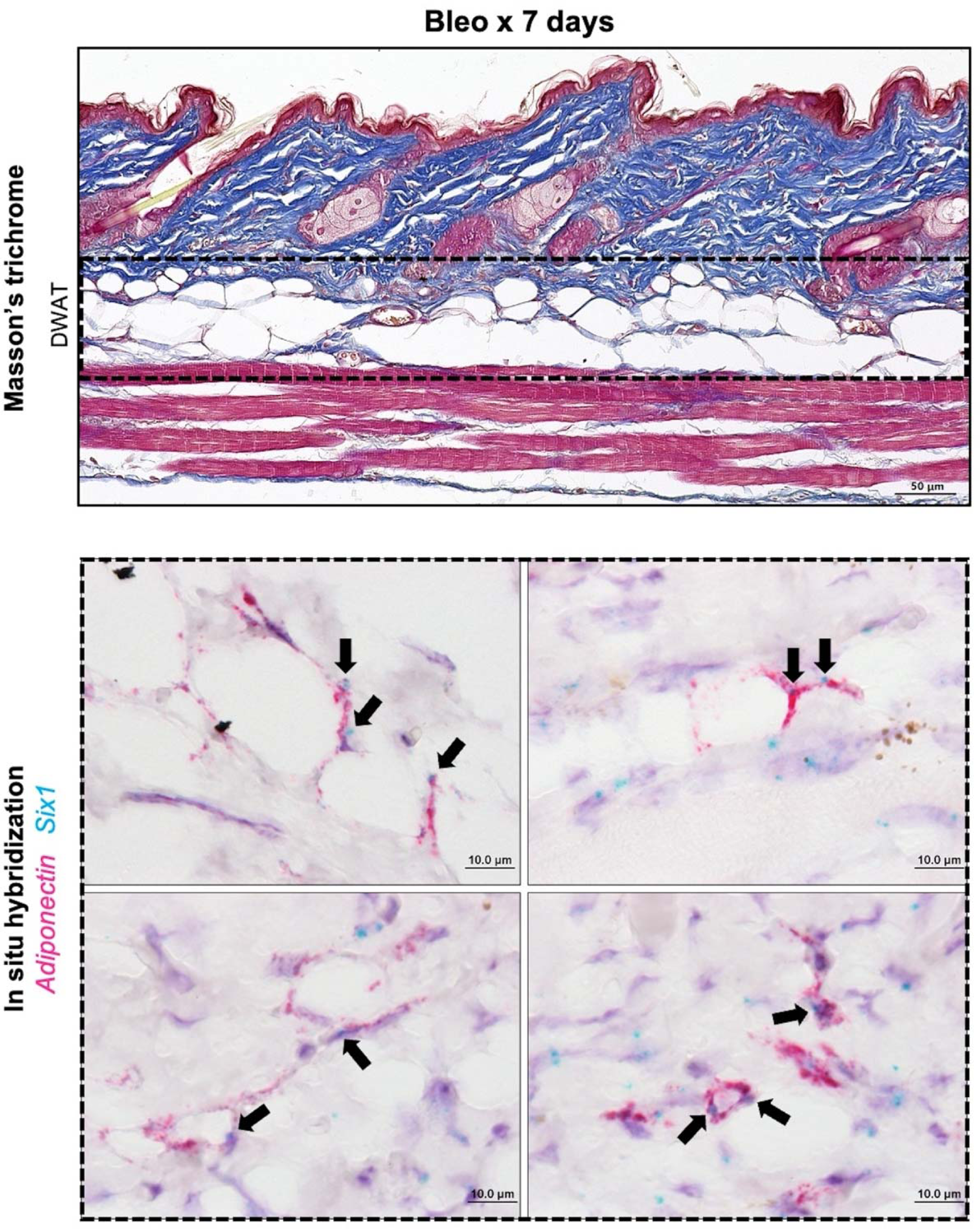
Adipocyte *Six1* expression precedes white adipose loss in SQ bleomycin-treated mice. Representative images of mouse dorsal skin injected with SQ bleo for 7 days (n=6). Top: Masson’s trichrome staining. Bottom: Dual *in situ* hybridization for *Adiponectin* (pink) and *Six1* (teal). Arrows point to *Six1* signal. DWAT=dermal white adipose tissue.

### Transgenic *Six1* deletion attenuates bleomycin-induced skin fibrosis

We next investigated whether genetic inhibition of *Six1* could prevent skin fibrosis. After a 28-day course of SQ bleo to induce an end-stage fibrosis phenotype, the affected skin of tamoxifen-treated mice with (iUbc^Cre^) and without the *Six1* allele (iUbc-Six1^-/-^) was analyzed by gene expression profiling and histology. *Six1* depletion in iUbc-Six1^-/-^ skin following tamoxifen was confirmed by RT-qPCR (**Figure 5 b**). Expression of profibrotic agents *Col1a1, Col1a2, Fn1, elastin, Acta2, Tgfb1 and Serpine1 but not Mif* was decreased in the skin of iUbc-Six1^-/-^ compared to iUbc^Cre^ mice (<u>**Figure 5 c-j**</u>). However, we did not detect changes in expression of adipocyte markers*: Adiponectin, Cebpa, or Pparg* (**Figure 5 k-m**). Masson’s trichrome staining of bleo-affected skin showed a protection of the DWAT layer in iUbc-Six1^-/-^ mice compared to iUbc^Cre^ mice (<u>**Figure 6a**</u>). *Six1*-deficient mice had more prominent DWAT with lipid-laden adipocytes (**Figure 6b).**

**Figure 5.**
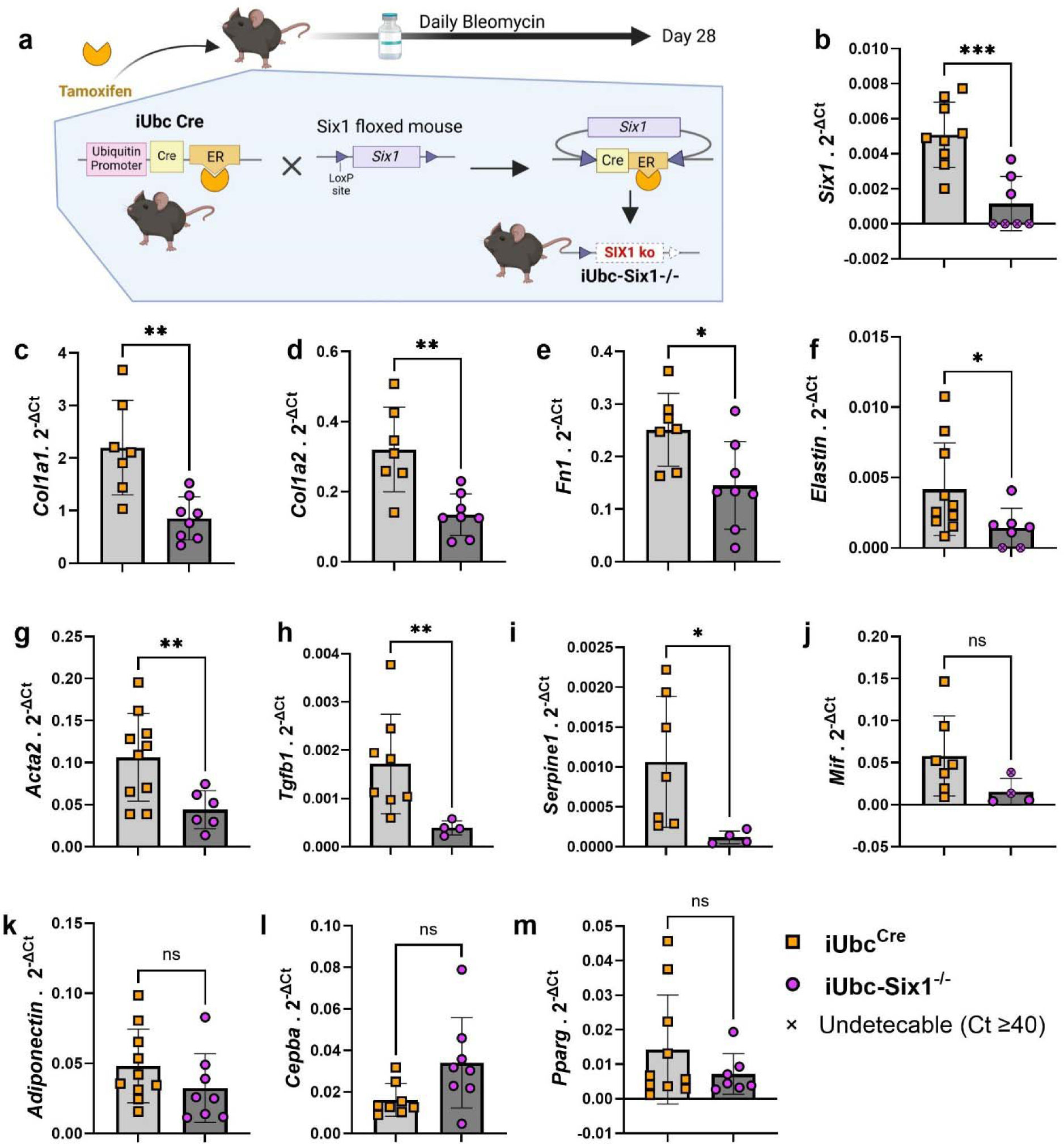
Inducible global deletion of Six1 inhibits fibrotic gene expression in SQ bleomycin treated mice. **a)** Schematic representation of experimental design (Created with BioRender). Following IP tamoxifen administration, iUbc^Cre^ and iUbc-Six1^-/-^ were given 28 days of SQ vehicle (PBS) or bleo. Transcript expression levels for sine oculis homeobox homolog 1 (*Six1,* **b**),collagen 1a1 (*Col1a1,* **c**), collagen 1a2 (*Col1a2,* **d**), fibronectin (*Fn1,* **e**), elastin (**f**), actin alpha 2 (*Acta2*, **g**), transforming growth factor beta 1 (*Tgfb1,* **h**), serpin family E member 1 (*Serpine1*, **i**), macrophage migration inhibitory factor (*Mif,* **j**), *Adiponectin* (**k**), CCAAT enhancer binding protein alpha (*Cebpa,* **l**), peroxisome proliferator activated receptor gamma (*Pparg,* **m**), at day 28 of bleomycin treatment.. Expression was normalized to 18s rRNA.. Significance levels * P ≤0.05, ** P ≤0.01, *** P ≤0.001, and **** P ≤0.0001 refer to an unpaired t-test for panels b-m. Each individual plot represents a biological N number.

**Figure 6.**
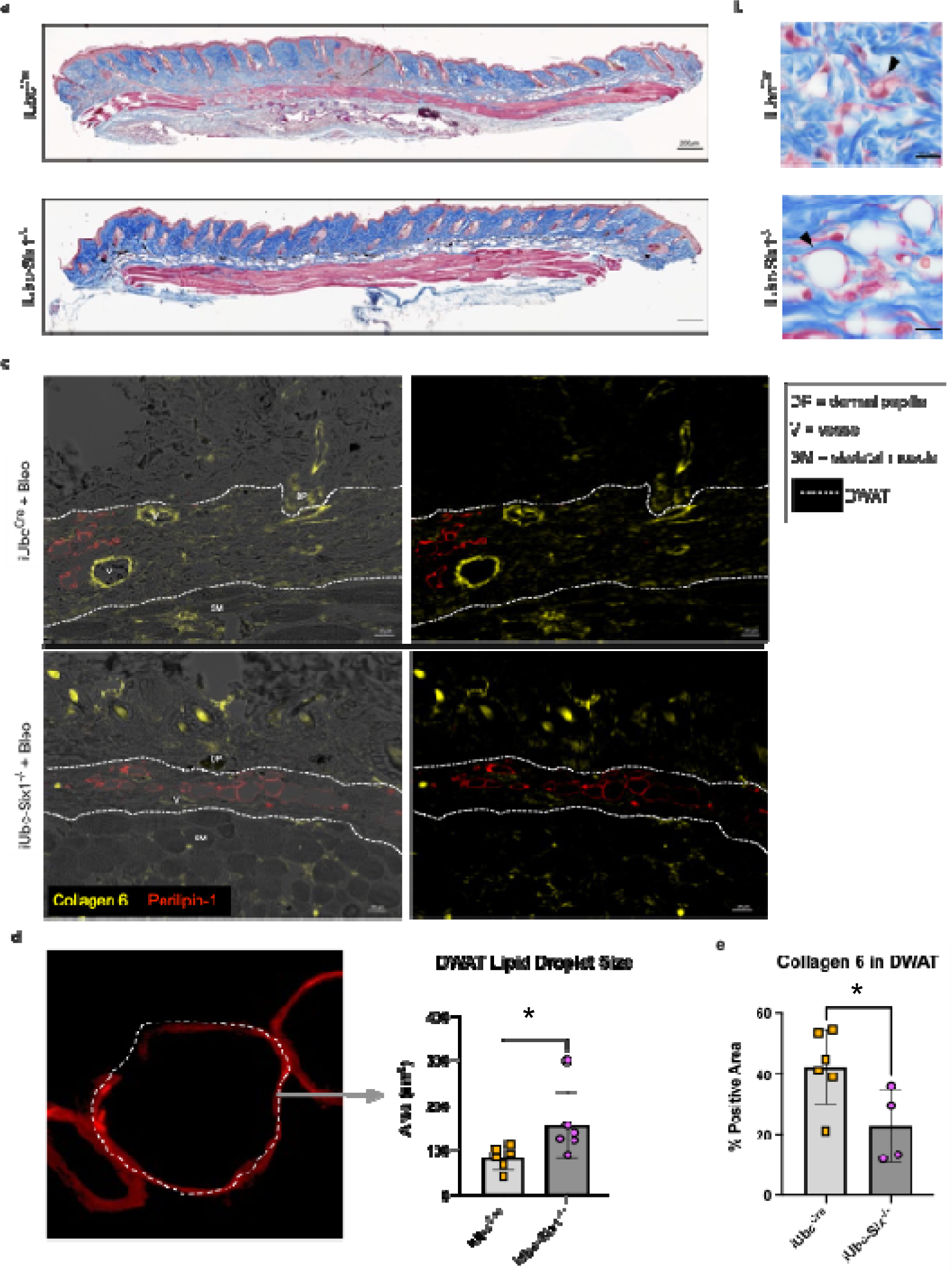
Global *Six1*-deletion prevents lipolysis and collagen 6 accumulation induced by SQ bleomycin. **a)** Masson’s trichrome staining of skin biopsies from iUbcCre and iUbc-Six1^-/-^ mice treated with 28 days of SQ bleo. **b)** High-magnification Masson’s trichrome images of DWAT. Black arrows point to lipid-laden adipocytes. **c)** Dual immunofluorescent staining for collagen 6 and perilipin 1 in skin from 28-day bleo-injected iUbc^Cre^ and iUbc-Six1^-/-^ mice. Left: Overlay of immunofluorescence images with bright-field images. Right: Immunofluorescence for perilipin 1 only. **d)** Quantification of droplet size of perilipin 1-positive adipocytes in the DWAT. Left: High magnification (100X, zoomed) image of lipid droplet within DWAT stained with perilipin 1. **e)** Quantification of collagen 6 as percent positive area in the DWAT. Significance levels * P ≤0.05 refer to an unpaired t-test. Each individual plot represents a biological N number.

Perilipin 1 immunostaining was used to specifically detect adipocyte lipid droplets (**Figure 6c).** Adipocyte droplets in iUbc-Six1^-/-^ were significantly larger compared to iUbc^Cre^ mice (**Figure 6d).** The deposition of collagen 6, which is enriched in adipose tissue, was analyzed using dual immunofluorescent staining with perilipin 1 to identify the DWAT (**Figure 6c**).^71,72^ iUbc-Six1^-/-^ had lower collagen 6 density in the DWAT compared to iUbc^Cre^ mice (**Figure 6e**). There was no significant difference in collagen 6 deposition in the dermis, or in dermal thickening (data not shown). In summary, whole-body depletion of *Six1* followed by 28 days of SQ bleo revealed that *Six1* deletion may halt pro-fibrotic gene expression and maintain DWAT in skin fibrosis.

### Adipocyte *Six1* expression drives dermal fibrosis

To build on our data demonstrating elevated *SIX1* in biopsies from individuals with early SSc and adipose *Six1* expression in a mouse model of dermal fibrosis, when DWAT has begun to atrophy. We next investigate the potential role of adipocyte specific Six1 in early disease. We developed a transgenic mouse model to knock-out *Six1* in cells with an active *Adipoq* promoter after tamoxifen treatment, conferring adipocyte-specific *Six1* depletion in adult mice (<u>**Figure 7a**</u>).^63^ We challenged mice with SQ vehicle or bleo for 14 days, as we found this duration of bleo induced DWAT atrophy before late fibrosis is established. Gene expression and histological analyses were performed to determine the contribution of adipocyte *Six1* in early events in skin fibrosis. Masson’s trichrome staining and dermal thickness measurements revealed a thinner dermis in iAdipo-Six1^-/-^ mice after SQ bleo compared to iAdipo^Cre^ mice (**Figure 7b-c**), demonstrating how adipocyte-specific *Six1* deletion prevented dermal thickening compared to *Six1*-competent mice. Next, we performed perilipin 1 immunostaining to identify adipocytes in the DWAT and measured intracellular lipid droplet size (<u>**Figure 8a**</u>). No difference in droplet size was observed in vehicle-treated mice regardless of Six1 expression. However, adipocyte SIX1 deletion inhibited the reduction in droplet size induced by bleo that precedes loss of adipose tissue (**Figure 8b**). These findings suggest that *Six1* plays a role in dermal lipid droplet size during the early development of skin fibrosis. To determine whether adipocyte preservation was accompanied by reduced DWAT collagen deposition, we performed immunofluorescent staining to detect collagen 6. While collagens 1, 4, and 6 are abundantly expressed in adipocytes, collagen 6 is the most predominant collagen in fat depots ^72^, thus the measurement of such would be expected to have the highest sensitivity for significant changes in the ECM. Samples were co-stained with perilipin 1 to identify the DWAT (<u>**Figure 9a**</u>). Deletion of *Six1* in adipocytes significantly reduced DWAT collagen 6 deposition after 14 days of bleo (**Figure 9b**). Adipocyte lipid droplet shrinkage and increased ECM deposition cause dramatic changes to DWAT architecture and function in skin fibrosis. We have here established adipocyte *Six1* as a driver of both lipid droplet size and ECM deposition, and a candidate target for therapeutic intervention.

**Figure 7.**
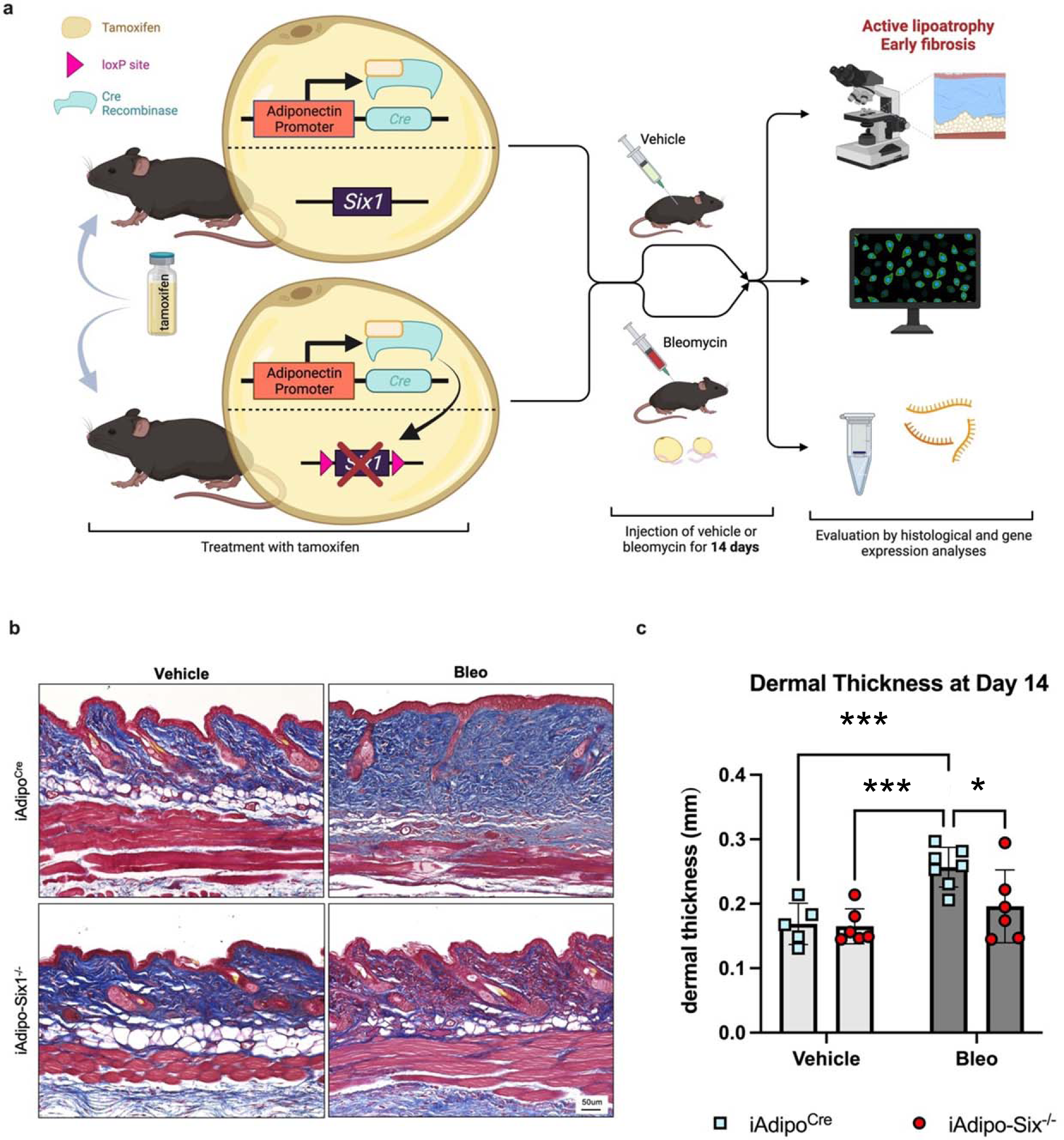
Adipocyte SIX1 deletion prevented bleomycin-induced dermal white adipose atrophy and dermal thickness. **a)** Schematic representation of experimental design (Created with BioRender). Following IP tamoxifen administration, iAdipo^Cre^ and iAdipo-Six1^-/-^ were given 14 days of SQ vehicle (PBS) or bleo. Full-thickness skin biopsies were harvested for histology and RNA. **b)** Representative images of Masson’s trichrome staining. **c)** Quantification of dermal thickness from Masson’s trichrome staining (Panel b). Significance levels * P ≤0.05 and ***P≤0.001 refer to a One-way ANOVA with a Bonferroni correction for panel c. Each individual plot represents a biological N number.

**Figure 8.**
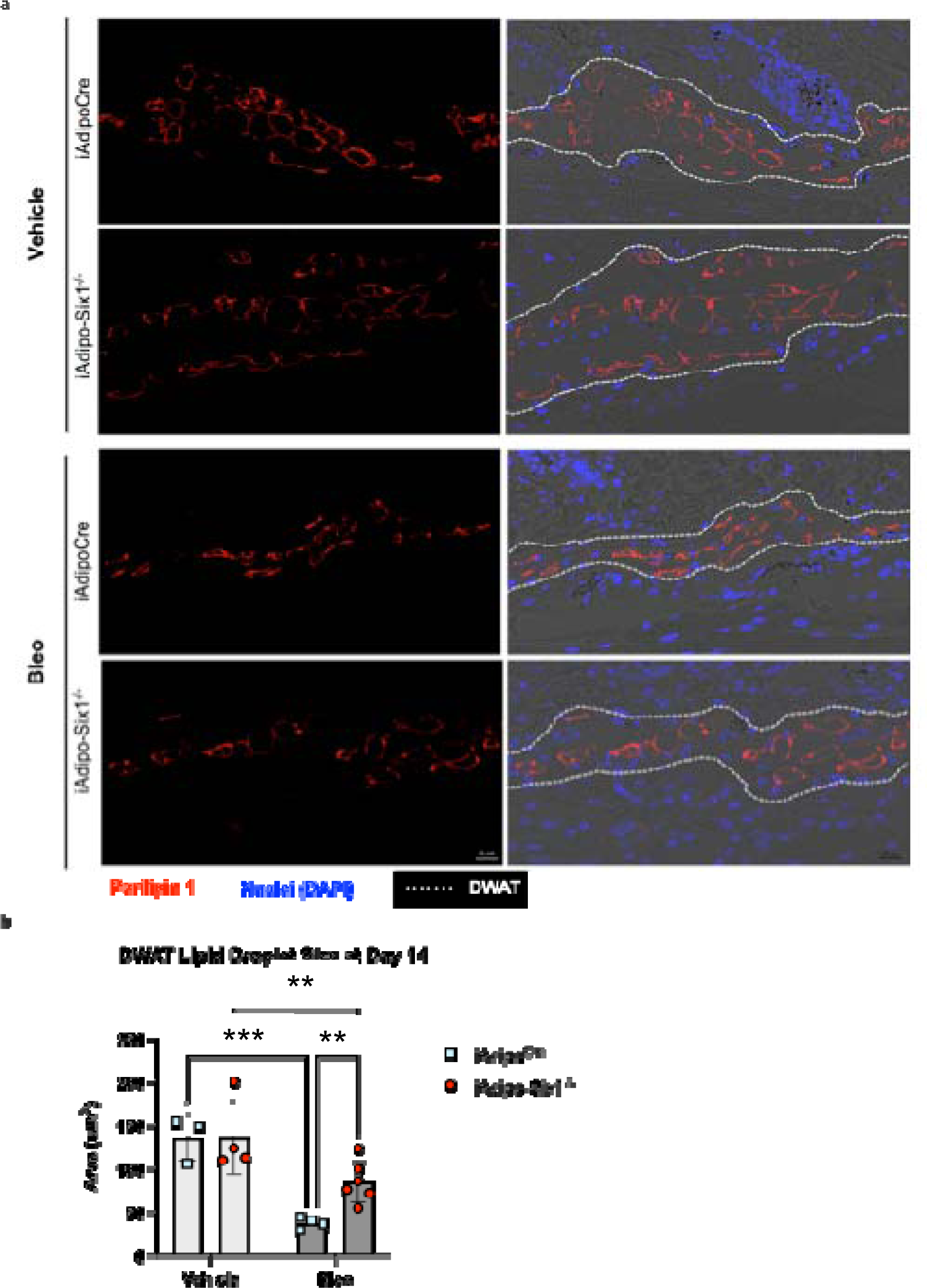
Lipid droplets are preserved in bleomycin-treated adipocyte-Six1 deficient mice trated with bleomycin. **a)** Representative images of immunofluorescent staining for perilipin 1 in skin from iAdipoc^Cre^ and iAdipo-Six1^-/-^ mice treated with 14 days of SQ vehicle (PBS) or bleo. Left: Immunofluorescence for perilipin 1 only. Right: Overlay of immunofluorescence images with bright-field images. **b)** Quantification of droplet size of perilipin-positive adipocytes in the DWAT. Significance levels ** P ≤0.01 and ***P≤0.001 refer to a Two-way ANOVA with multiple comparison employing a Sidak correction for panel b. Each individual plot represents a biological N number

**Figure 9.**
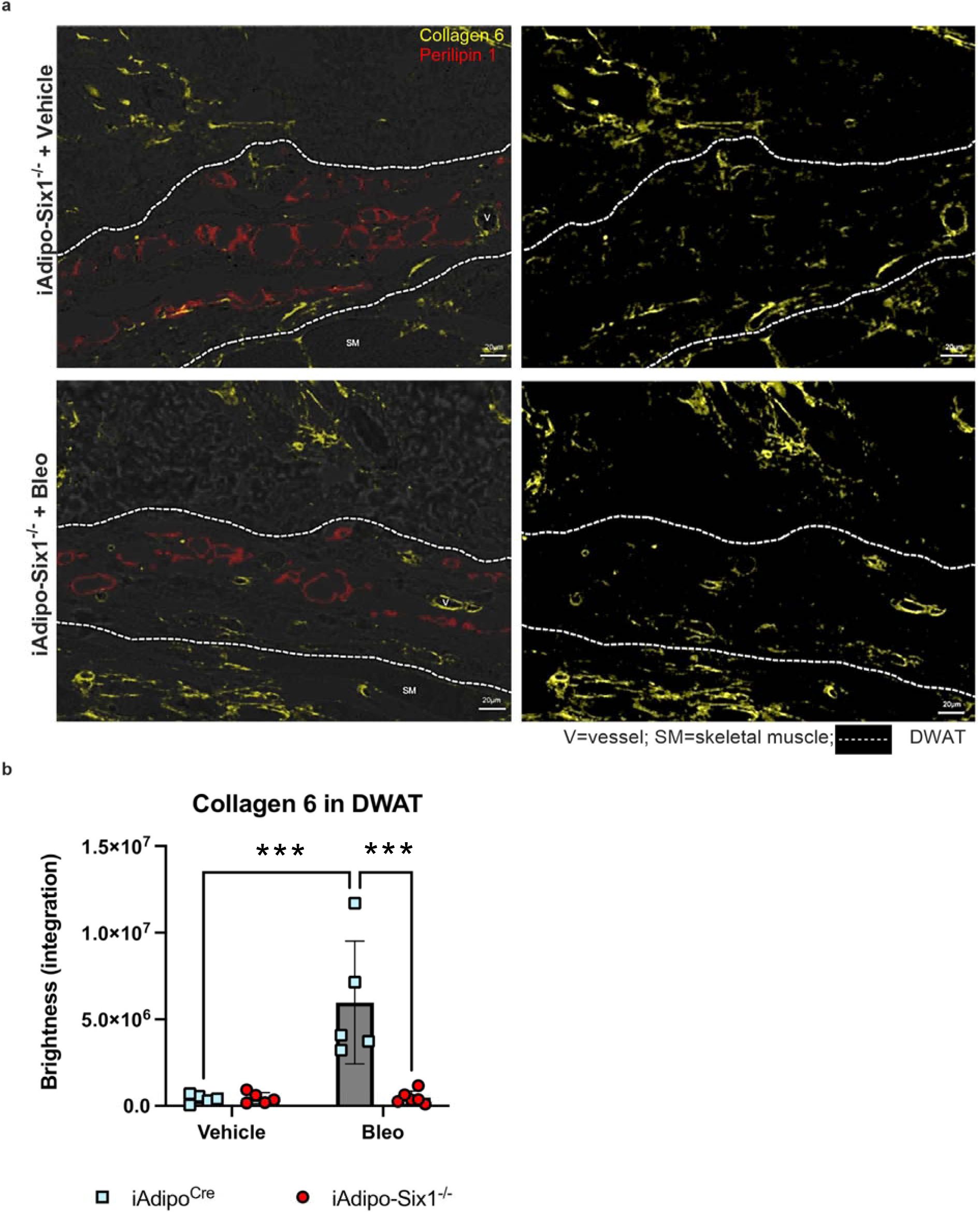
iAdipo^Cre^ -*Six1* KO mice have decreased collagen 6 deposition in the DWAT after 14 days of SQ bleomycin. **a)** Representative images of dual immunofluorescent staining for collagen 6 (yellow) and perilipin 1 (red) in skin samples from iAdipo-Six1-/- mice treated with 14 days of SQ vehicle (PBS) or bleo. *Left*: Overlay of immunofluorescence with bright-field images. *Right*: Immunofluorescence for collagen 6 only. Area within dotted lines represents DWAT as identified by positive perilipin 1 staining. **b)**. Quantification of collagen 6 as brightness intensity within the DWAT. Significance levels ***P≤0.001 refer to a Two-way ANOVA with multiple comparison employing a Sidak correction for panel b. Each individual plot represents a biological N number

Collectively, our data demonstrates that SIX1 deletion helps maintain lipid droplet size and ECM deposition in bleomycin treated mice. However, whether SIX1 can directly regulate stromal cell fate in dermal fibrosis is not known. Thus, to identify if SIX1 regulates cell fate, we used an *in vitro* cell differentiation model. We treated mouse 3T3L1 fibroblasts with siSIX1 (or control siRNA) and an adipocyte differentiation cocktail, to promote differentiation to adipose cells. Cells were collected on days 0, 2, 4, 6 and 9. These experiments demonstrated a successful SIX1 knockdown on day 4 and 6 by western blots (**Supplementary Figure 2 a, b**). Despite increased expression of adiponectin on days 4, 6 and 9 (**Supplementary Figure 2 c**) and Cebpa and Ppparg on days 6 and 9 (**Supplementary Figure 2 d, e**), reduced levels of SIX1 did not alter the expression of these lipid mediators, with the exception of Cebpa on day 9. Similarly, although we report an increase of *Tgfb1* on days 6 and 9 of the differentiation cocktail, SIX1 KD did not alter expression levels of *Tgfb1* (**Supplementary Figure 2 f**). In line with inability of SIX1KD to alter lipid expression levels, we report no changes in the differentiation of 3T3L1 cells to adipocytes as seen by oil-red O staining (**Supplementary Figure 2 g**).

Next, we performed a gene analysis using the nCounter platform targeting fibrotic gene expression on Day 6 of 3T3 treated fibroblasts with and without siSIX1. These unbiased experiments and subsequent heat maps for ECM synthesis and TGF-β signaling revealed that KD of SIX1 reduced levels of serine proteinase inhibitor E 1 (SERPINE1) the gene that encodes plasminogen activator inhibitor 1 (PAI-1), (**Supplementary Figure 3a, b)**. These results also demonstrated reduced Col4a1, Col4a2, Col5a1 and Col5a3 and Col6a3 in addition to Tgfb1. Analysis of the TGF-β signaling pathway revealed reduced downstream mediators following siSix1 such as Tgfbr1, Crebbp, or Furin (**Supplementary Figure 3a, b**). Serpine1 was selected for further validation based on a volcano plot demonstrating that it was one of the most significantly downregulated genes following siSix1. (**Supplementary Figure 4**). These findings were confirmed both by qPCR and Western blots and corresponding densitometries for Serpine1 and PAI-1 respectively from 3T3 cells, revealing increased Serpine1 transcript and protein expression levels on D9 that were inhibited following siSIX1 treatment (**Figure 10 a, b**). These studies were consistent with IHC from SQ bleo treated mice demonstrating increased expression levels of PAI-1 in DWAT region that was inhibited in mice with deleted *Six1* in adipocytes (**Figure 10c**). Expression of SERPINE1 was also upregulated 11.5-fold and 3.8-fold in the PRESS and GENISOS cohorts respectively (**Table 1**). Although the increased expression of SERPINE-1 could This suggest that these mediators may be involved in promoting the dermal profibrotic response in SSc, yet further experiments are necessary to identify increased expression This points at elevated *SERPINE1* and its protein PAI-1 as a downstream pro-fibrotic mediator of *SIX1* upregulation in adipocytes as a potential mechanism leading to dermal fibrosis.

**Figure 10.**
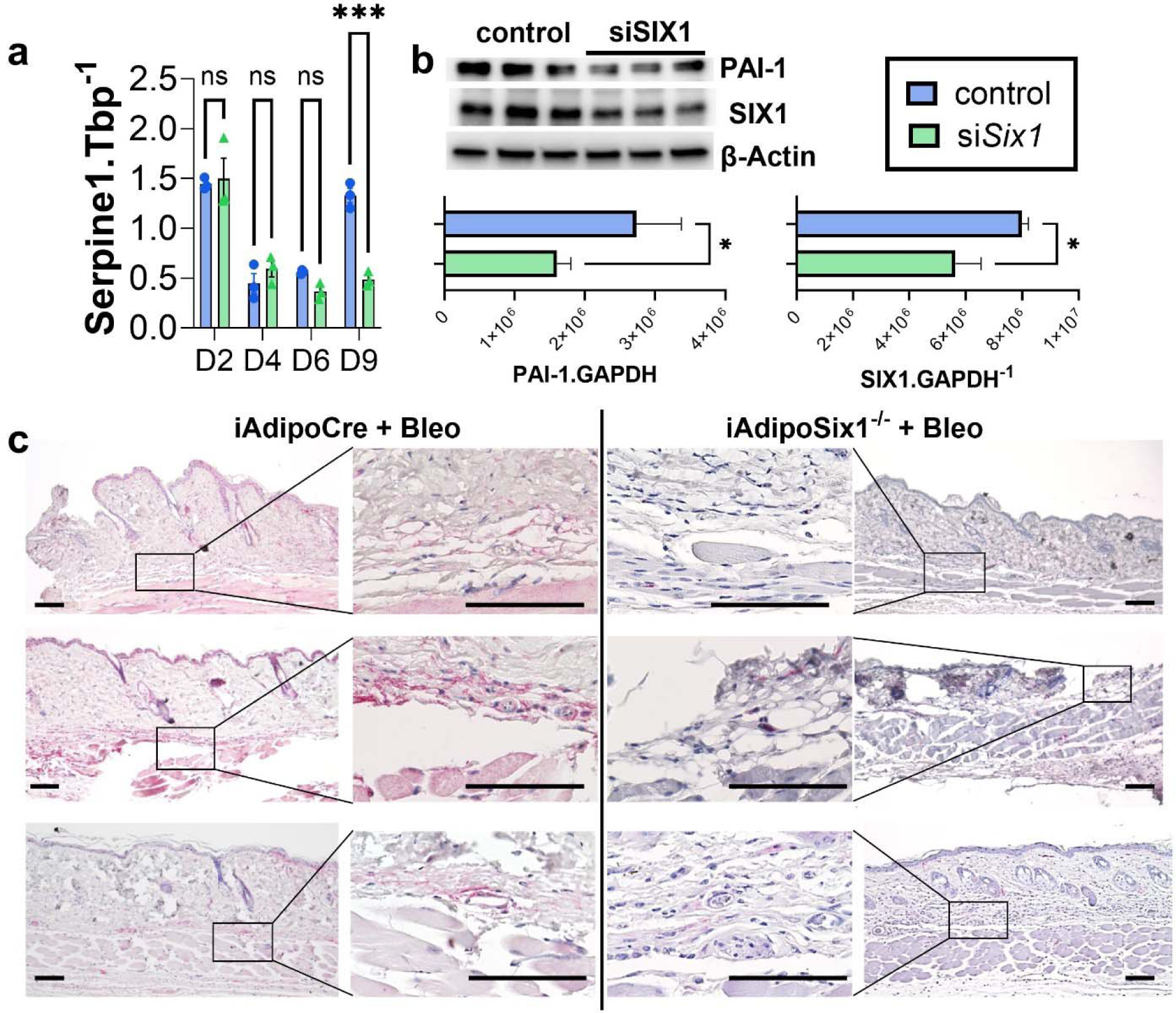
PAI-1 levels are upregulated in bleomycin induced skin fibrosis and track with SIX1 expression levels. **a)** Serpine1 transcript levels and (**b**) protein levels for PAI-1, SIX1 and β-actin from 3T3 cells treated with a differentiation cocktail to adipocytes and transfected with either control siRNA (blue bars, panel a) or siSix1 (green bars, panel a). Densitometries for PAI-1 and SIX1 expression relative to GAPDH is shown below. **c**) IHC for PAI-1 from bleomycin exposed mouse skin for from 3 independent iAdipoCre or iAdipoSix1^-/-^ mice. PAI-1 signals are shown in magenta. Significance levels ***P≤0.001 refer to a Two-way ANOVA with multiple comparison employing a Sidak correction for panel a. Significance levels *P≤0.05 refer to an unpaired t-test for panel b. N=3 for all data.

**Table 1.** SERPINE1 expression levels in GENISOS and PRESS cohorts.

| **Cohort** | **GENISOS** | **PRESS** |
| --- | --- | --- |
| Ensembl_ID | ENSG00000106366.8 |  |
| Gene name | SERPINE1 |  |
| Gene type | Protein coding |  |
| logFC | 1.928211 | 3.524306 |
| logCPM | 2.589225 | 4.593314 |
| **PValue** | **2.64E-12** | **4.17E-33** |
| FDR | 2.05E-09 | 9.2E-30 |
| **FC** | **3.805831** | **11.50593** |
| CPM | 6.017755 | 24.13933 |
| Regulation in SSc group | Up | up |

FC: Fold Change, CPM: Counts per million

## Discussion

This study demonstrates the previously undescribed clinical and translational relevance of the developmental transcription factor *SIX1* in adipocyte mediated skin fibrosis. We identified increased expression of the developmental transcription factor *SIX1* in skin-associated adipose tissue in SSc skin samples from two well-described cohorts encompassing 161 patients. The GENISOS cohort contains both lcSSc and dcSSc, allowing us to identify the further increase in *SIX1* in diffuse disease in a large cohort. *SIX1* was also elevated in the skin of individuals with dcSSc diagnosed within three years of disease onset, obtained from the PRESS cohort. In agreement with expression data showing enrichment of *SIX1* in dcSSc, *SIX1* skin expression positively correlated with clinical measurements consistent with worse skin disease extent and severity. Previous work established a subcutaneous adipose signature in human skin.^51,52,61^ We found this signature to be strongly correlated with *SIX1* expression in both SSc cohorts. It is important to note that a specific dermal adipose signature has not yet been established.^69^ This observation suggests that in SSc skin, *SIX1* expression may be associated with genomic changes in adipocyte function. Expression data was supported by histology findings in SSc skin samples from the GENISOS cohort. We reproduced previously reported observations^15,65^ that dermal fat mass declines early and remains atrophic in SSc skin. Using two novel transgenic mouse models, we found evidence for a role for *Six1* in lipodystrophy and fibrotic features observed in the SQ bleo model. Intriguingly however, only deletion of *Six1* from adipocytes but not in UBC^Cre^ expressing mice resulted in a reduction of dermal thickening. Possible explanations for this include a more efficient and selective inhibition of *Six1* in adipocytes through the adiponectin Cre system compared to UBC^Cre^ expression or a potential protective response of *Six1* in other cells that could include mesenchymal or inflammatory cells. We propose a working model in which SIX1 helps to drive skin fibrosis by interacting with lipolysis-associated molecular pathways to promote intracellular lipid loss, a critical first step towards transforming a healthy adipocyte to a disease-driving cell type. However, it is also possible for the adipocyte layer to function as a protective element against the development of fibrosis that is lost as tissue atrophies.

We found that *SIX1* expression in SSc skin correlated with genes related to lipid metabolism. We and others have shown that lipid droplets in DWAT are smaller in the SQ bleo-treated mice.^22,73^ Mice with *Six1-*deficient adipocytes (*Six1*^-^ WAs) had larger intracellular lipid droplets than *Six1*^+^ WAs after SQ bleo. Changes in size of unilocular lipid droplets in WAs is one surrogate measurement used to profile an adipose depot as being lipogenic or lipoatrophic.^66,74^ A critical step in AMT is the release of free fatty acids into the local tissue environment, a process which permits the transition of a lipid droplet-containing adipocytes into a precursor cell^20^. AMT is a fluid process involving dynamic and complex changes in cellular phenotypes and gene expression^24,75^. Dynamic control of lipid storage is required for differentiation and transdifferentiation ^62,76,77^. We found *Six1* to be expressed in mouse DWAT after just 7 days of bleo. While the majority of *Six1*-positive (Six1^+^) cells were also positive for *Adipoq* (*Adipoq*^+^), we observed a minority of *Six1*^+^ *Adipoq*^-^ cells, that is, cells other than lipid-laden adipocytes. *Adipoq*^-^ cells in the stroma vascular fraction (SVF) include diverse cell types.^78^ Further*, ADIPOQ* was highly correlated with *SIX1* in SSc skin. Given the exclusivity of *Six1* expression to the DWAT, we propose that these cells most likely represent those of adipocyte lineage. These studies and the known capacity of SIX1 to promote trans-differentiation ^38,79^ point at a role for SIX1 in mediating AMT. Intriguingly, our fibroblast to adipocyte cell differentiation studies *in vitro*, revealed that inhibition of Six1 in 3T3L1 cells did not prevent fibroblasts becoming adipocytes nor did it affect expression levels of adipocytes markers such as adiponectin, Cebp or Pparg. This was consistent with expression levels for these mediators that were unchanged in mice deficient in SIX1. However, using a non-biased approach using the nCounter platform, deletion of Six1 in 3T3 cells inhibited ECM components such as Col6a3, consistent with our IHC for Col6 and other collagens such as Col4a1, Col4a2, Col5a1 and Col5a3 in addition to Tgfb1. Although *Tgfb1*^79^ and *Pparg*^43^ have been identified as targets of SIX1 our validation studies did not show significant changes following SIX1 KD, similar to *Mif* levels another target for SIX1^30^, highlighting the pleiotropic and cellular specificity of SIX1.

However, a common mediator that was altered in both heat maps was Serpine1, the gene encoding for PAI-1. These results revealed reduced Serpine1 following siSix1, a result that was validated by RT-qPCR and western blots and by IHC in skin section from bleo-treated mice revealing reduced PAI-1 signals in mice lacking Six1 in adipocytes. These results are significant since elevated PAI-1 has been shown to be elevated in skin lesions from SSc patients^80,81^ and its inhibition improved dermal inflammation and fibrosis in bleo treated mice.^81^ These findings suggest SIX1 as upstream from PAI-1 and as a mediator that predisposes adipocytes to a pro-fibrotic phenotype.

Targeted therapeutics in SSc are limited as a result of our fragmented understanding of disease mechanisms.^82^ Two medications, nintedanib and tocilizumab, have been approved by the Food and Drug Administration (FDA) for SSc-related interstitial lung disease.^83^ However, neither these drugs or others are FDA-approved for SSc skin involvement. SIX1 has been shown to be a potential therapeutic target in pulmonary fibrosis.^30^ We proposed that SIX1 might also play a role in skin fibrosis. We have shown that genetic deletion of *Six1* is able to reduce dermal adipose tissue atrophy and fibrotic changes in a rodent model of skin fibrosis. These studies demonstrate that adipose tissue homeostasis not only has anti-fibrotic effects, but that its preservation is a potential therapeutic approach for skin manifestations in SSc. Future studies should address whether pharmacological inhibition of SIX1 may serve such a purpose.

## Supporting information

Review comments

## Acknowledgments

We acknowledge PRESS and GENISOS investigators for their contribution to sample and data collection. Funding: Rheumatology Research Foundation Future Physician Scientist Award (Wareing); NIH (NHLBI) R01HL138510, R01HL157100 (Karmouty-Quintana); National Scleroderma Foundation Established Investigator Award (Karmouty-Quintana); NIH/NIAMS R01AR073284 (Assassi-Mills), NIH/NIAMS R01AR081280 (Assassi-Wu), NIH/NIAMS K08AR081402 (Skaug), Rheumatology Research Foundation Investigator Award (Skaug) NIH 1UL1TR003167 (W Jim Zheng) DoD W81XWH-22-1-0164 (W Jim Zheng). Susan Majka, PhD for critically reviewing the manuscript.

---

**Supplementary Figure 1.**
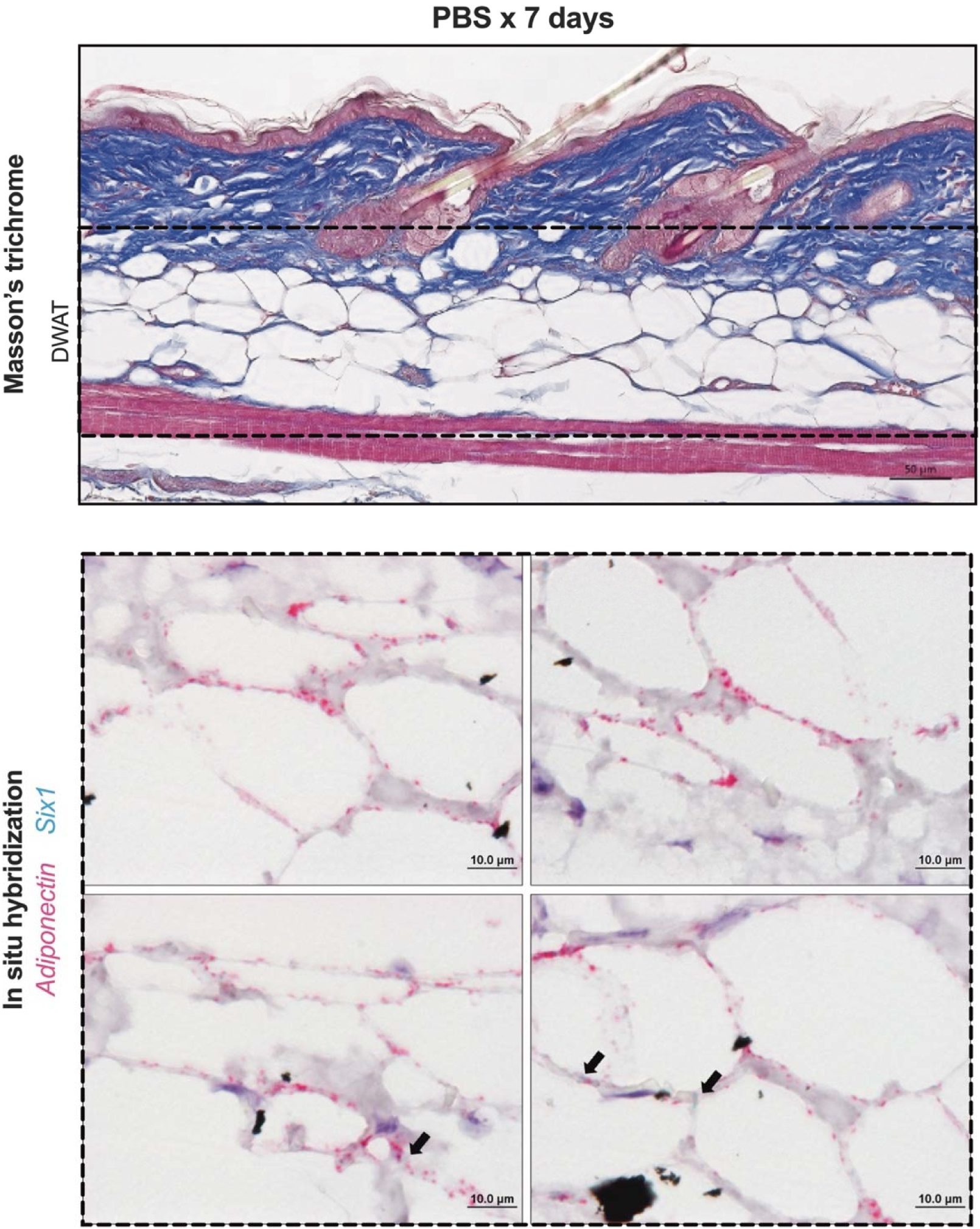
Six1 is minimally expressed in adipocytes in 7-day vehicle-treated mouse skin. Representative images of mouse dorsal skin injected with SQ vehicle (PBS) for 7 days (n=5). Top: Masson’s trichrome staining. DWAT=dermal white adipose tissue. Bottom: Dual *in situ* hybridization for *Adiponectin*/*Adipoq* (pink) and *Six1* (teal). Arrows point to *Six1* signal.

**Supplementary Figure 2.**
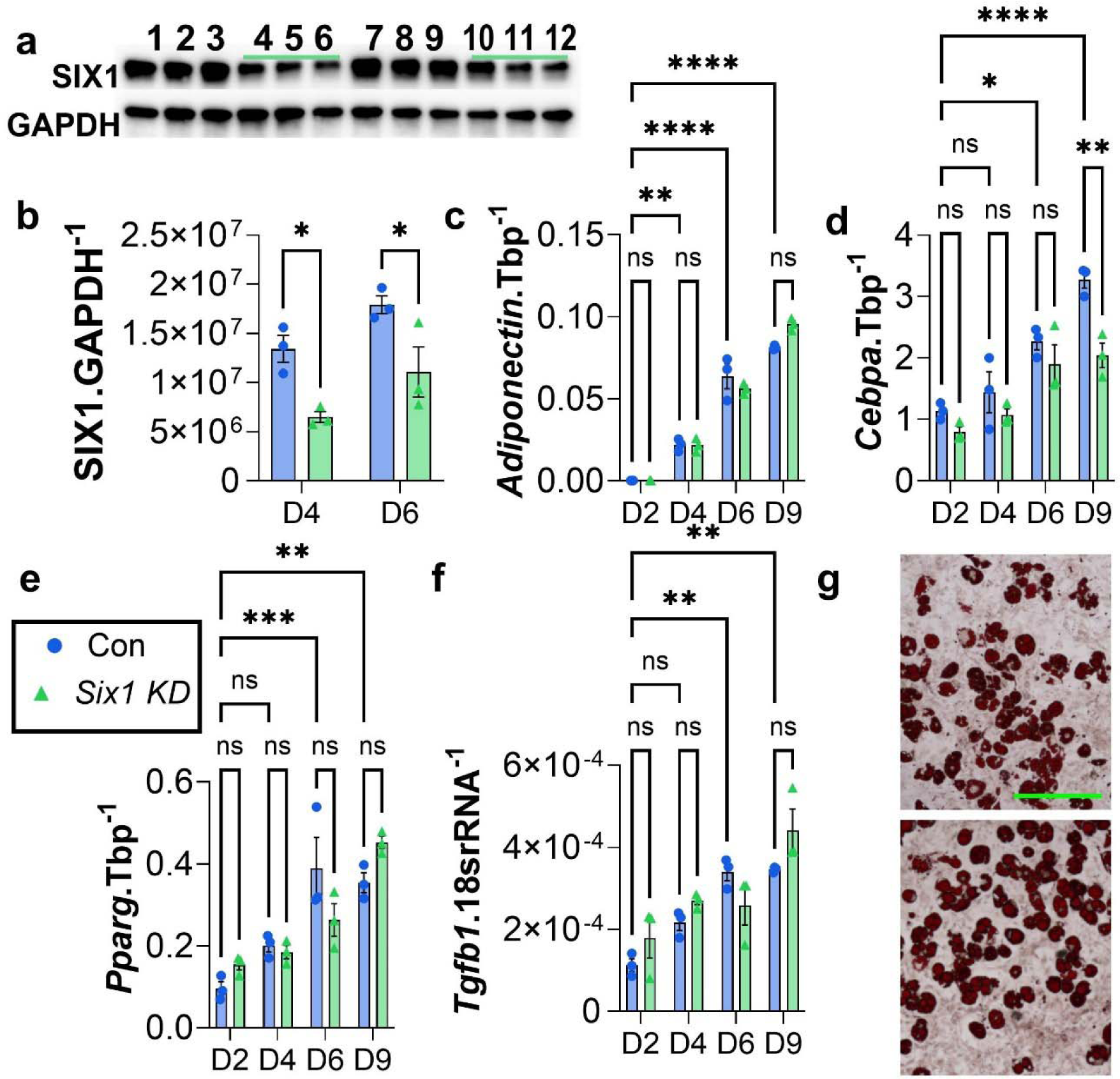
siSix1 transfection was effective at days 4 and 6 and minimally changed markers for adipocytes. 3T3 cells treated with a differentiation cocktail to adipocytes and transfected with either scrRNA (blue bars) or siSix1 (green bars). **a**) western blot for SIX1 or GAPDH lanes 1-3 represent scrRNA transfection on day 4, lanes 4-6 represent siSIX1 transfection on day 4, lanes 7-9 represent scrRNA transfection on day 6 and lanes 10-12 represent siSIX1 transfection on day 6.). Densitometries for SIX1 expression relative to GAPDH on days 4 and 6 (**a**). Expression levels for (**c**) adiponectin, (**d**) Cebpa (**e**) Pparg and (**f**) Tgfb1 from scRNA or siSix1 transfected 3T3 cells on days 2, 4, 6 and 9 after treatment with the differentiation cocktail. (**g**) Oil red O staining demonstrating formation of lipids from control (top) and SIX1 KD (bottom) 3T3L1 cells at day 9. Scale bar represents 50µm. Significance levels * P ≤0.05, ** P ≤0.01, *** P ≤0.001, and **** P ≤0.0001 refer to a Two-way ANOVA with multiple comparison employing a Sidak correction for panels b-f.

**Supplementary Figure 3.**
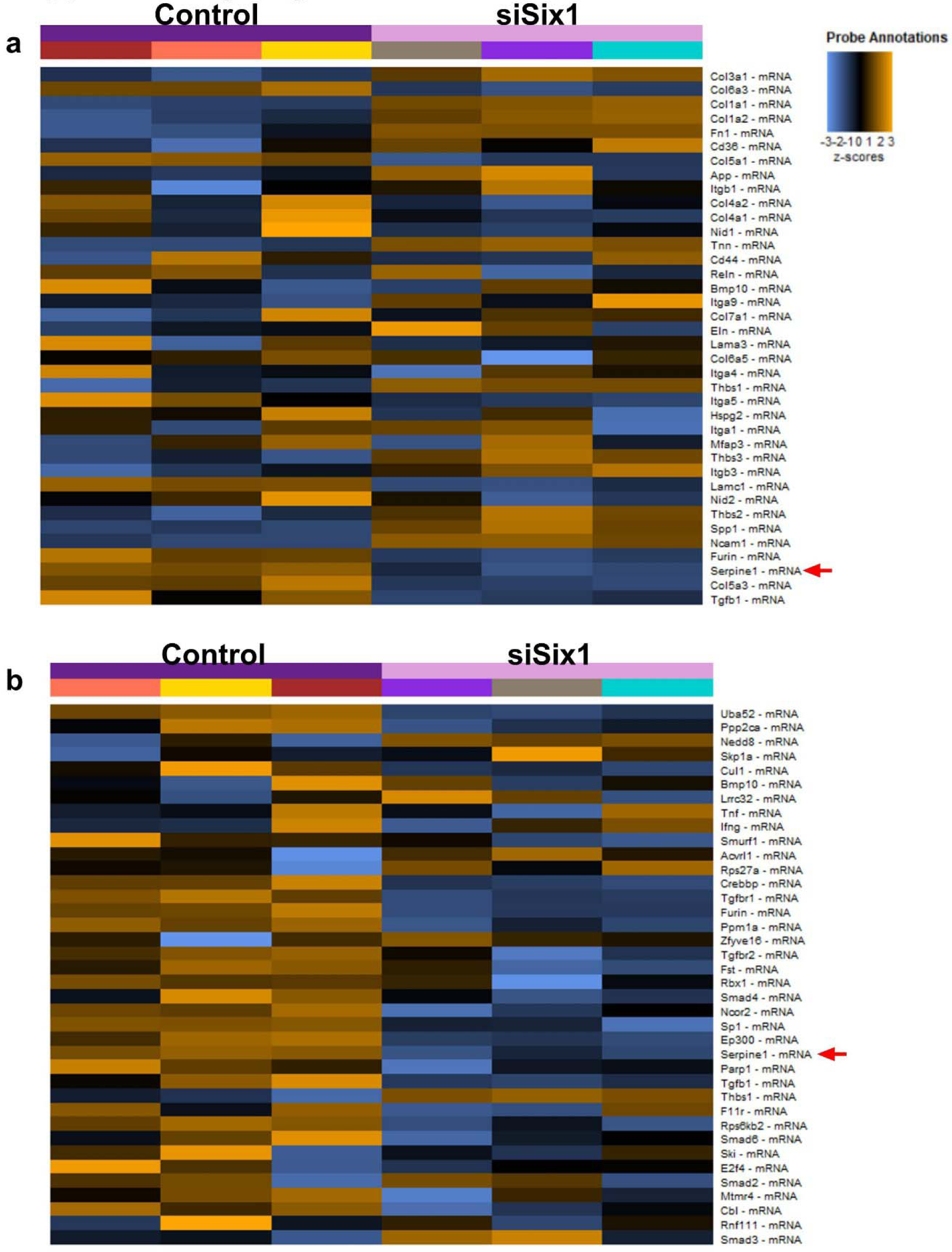
Heat maps from nCounter data identifying Serpine1 as a target. nCounter generated heatmaps for extracellular matrix (ECM) synthesis pathways (**a**) and TGF-β signaling pathway (**b**). Yellow shades represent upregulated genes and blue shades represent downregulated genes. The red arrow points at Serpine1.

**Supplementary Figure 4.**
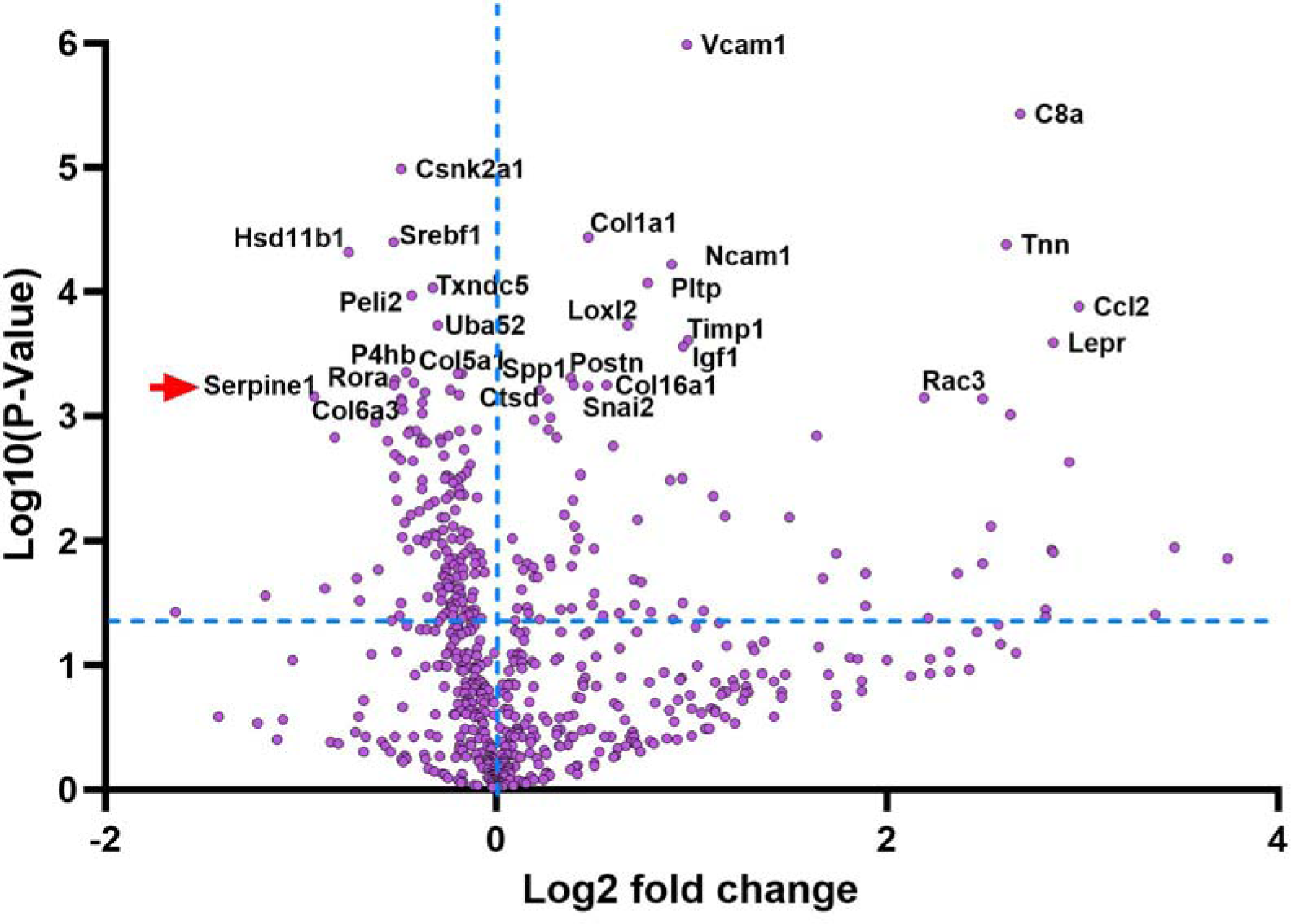
Volcano plot from nCounter data identifying Serpine1 as a target following SIX1 deletion. nCounter generated DEG volcano plot from Day 6 scRNA vs siSix1 transfected 3T3 cells treated with a differentiation cocktail to adipocytes. Significant genes are located above the horizontal dotted blue line. The top genes with altered expression are labelled including Serpine 1 identified by a red arrow.

**Supplementary Table 1:**
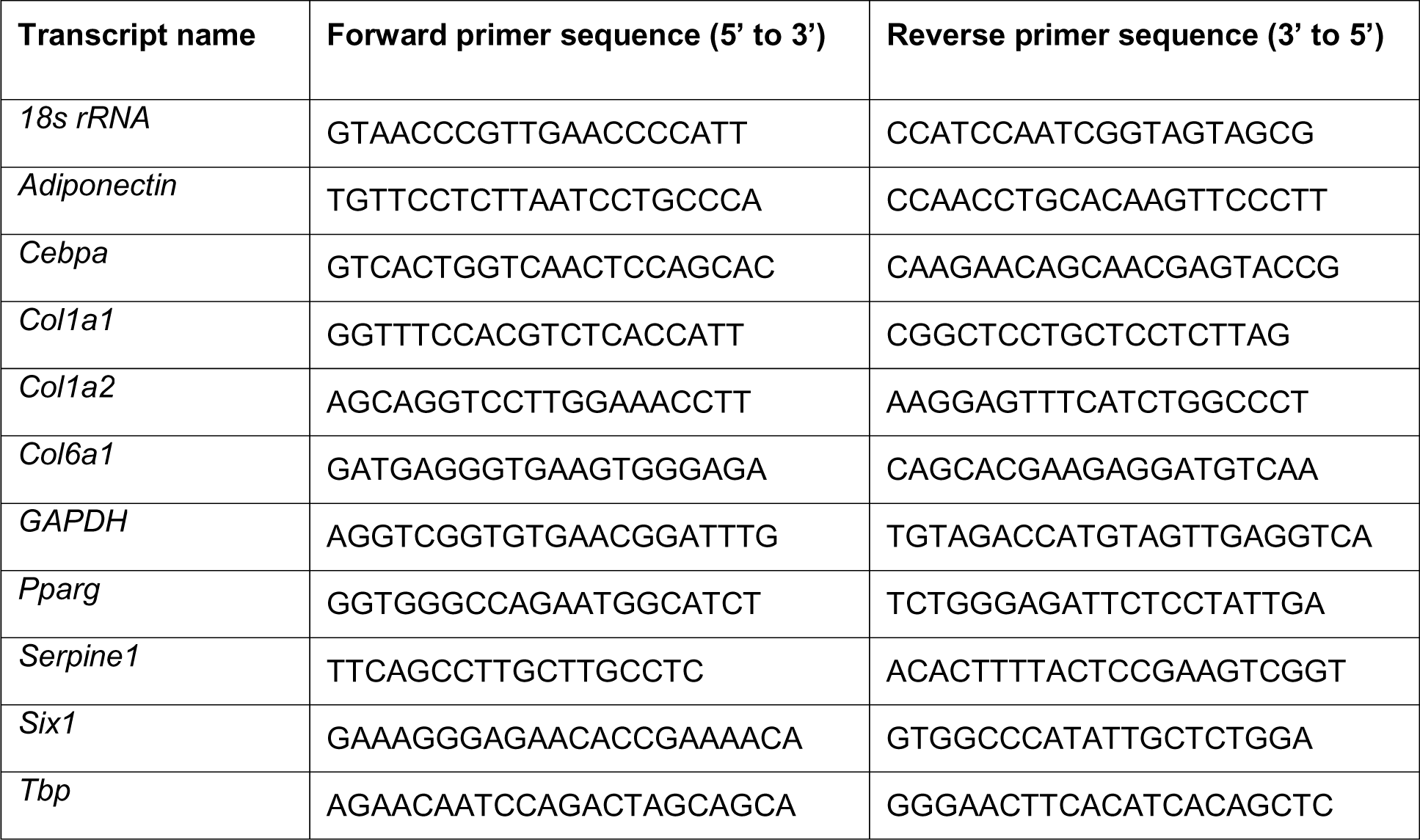
Mouse SYBR green primers used for quantitative polymerase chain reaction.

**Supplementary Table 2:** RNAScope probes used for in situ hybridization.

| **Target transcript (RefSeq Accession number)** | **Target species** | **Target region** | **Catalog #** |
| --- | --- | --- | --- |
| *SIX1* (NM_005982.3) | Human | 799-2177 | ACD; 412401 |
| *FABP4* (NM_001442.2) | Human | 119 - 826 | ACD; 470641-C2 |
| *Six1* (NM_009189.3) | Mouse | 1932-2819 | ACD; 838641 |
| *Adiponectin* (NM_009605.4) | Mouse | 9 - 1233 | ACD; 440051-C3 |

**Supplementary Table 3.**
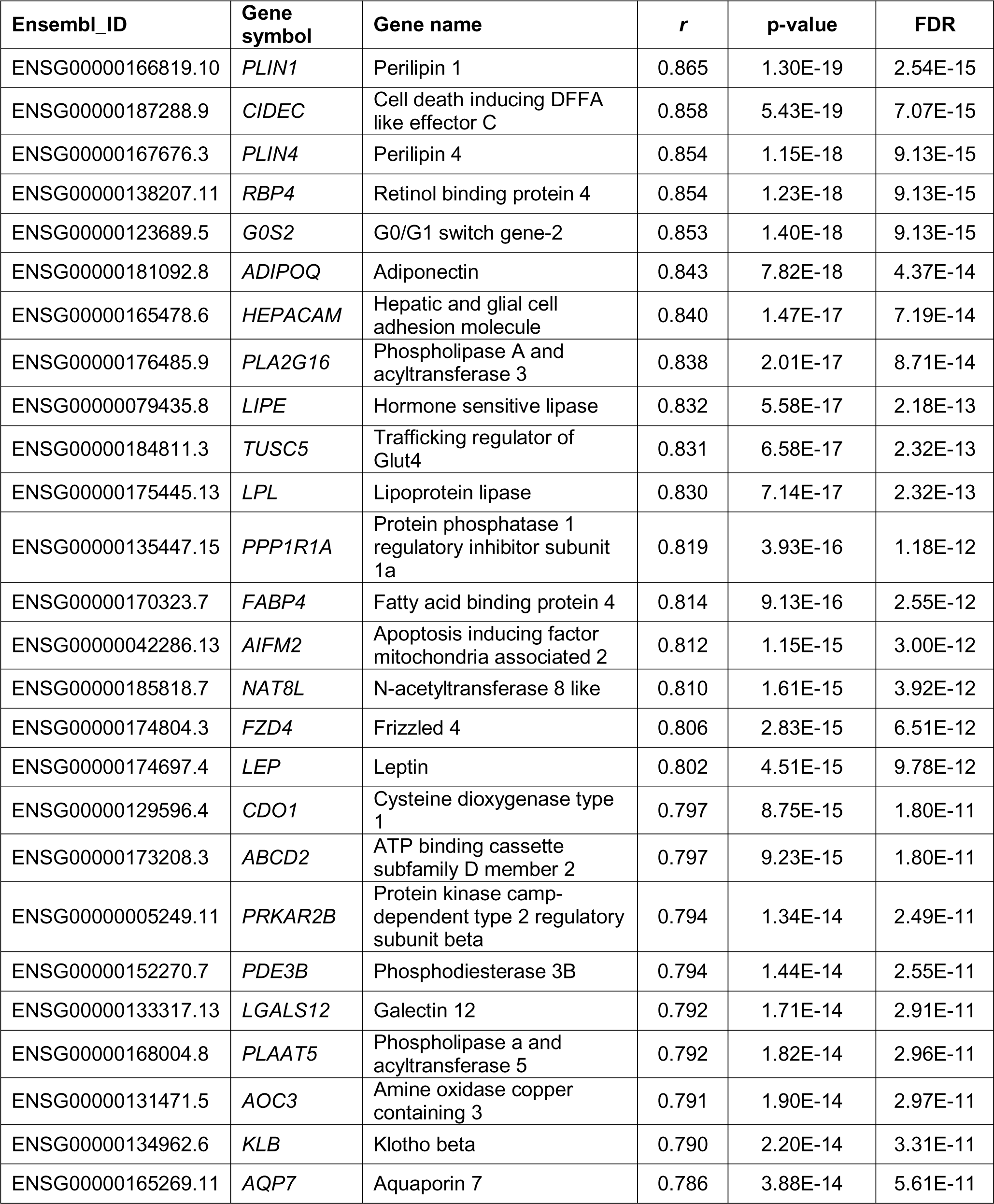
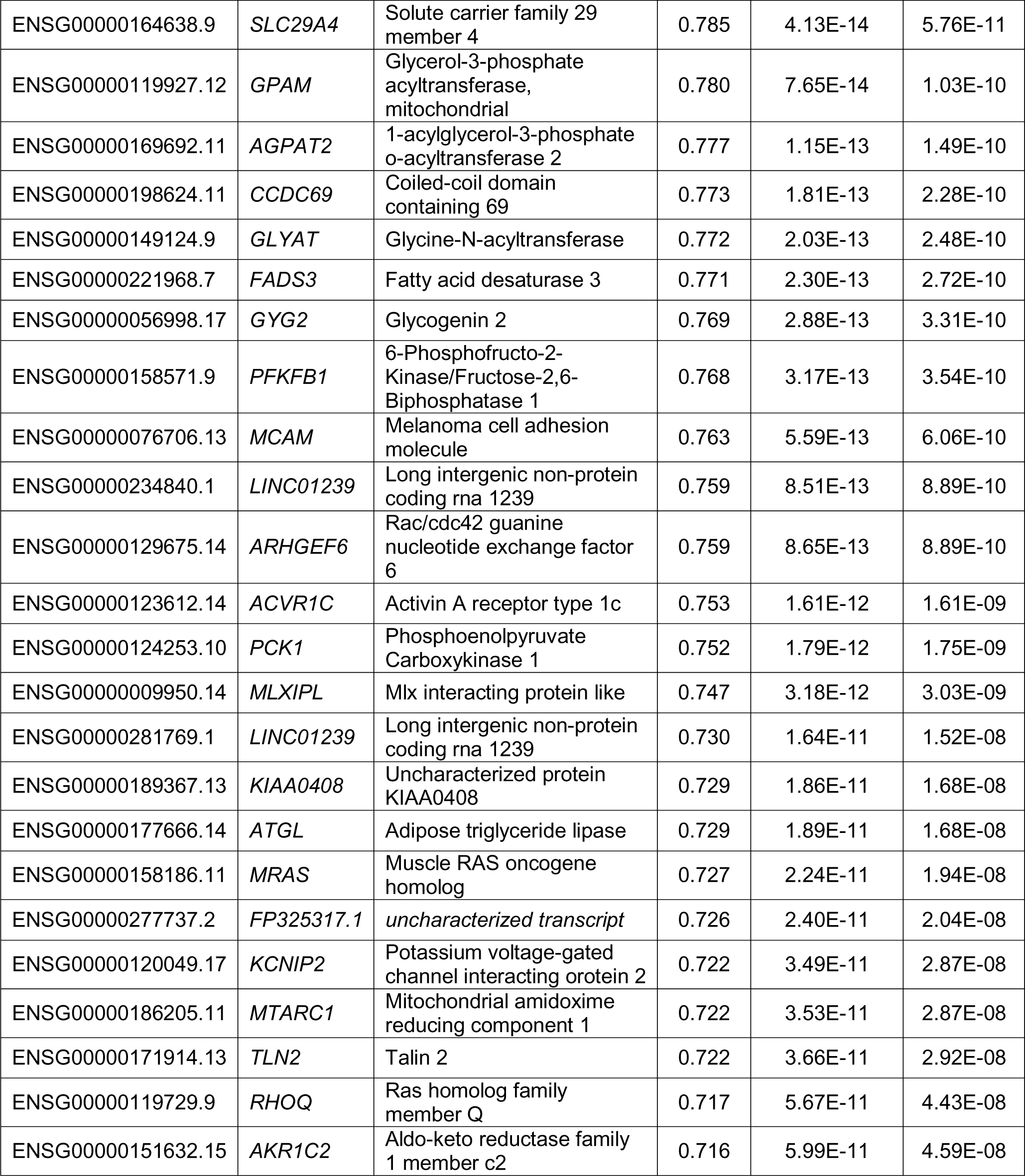
Top fifty most highly correlated genes with skin *SIX1* in the PRESS cohort by Spearman correlation analysis.

**Supplementary Table 4.** Demographic and clinical features of biopsied participants in Supplementary Figure 1.

| **Panel** | **Age (yrs)** | **Sex** | **Race** | **Disease subtype** | **Disease duration (yrs)** | **mRSS** | **local skin score** |
| --- | --- | --- | --- | --- | --- | --- | --- |
| a | 54 | Female | Caucasian | Control | N/A | N/A | N/A |
| b | 45 | Female | Caucasian | dcSSc | 2.3 | 22 | 2 |
| c | 62 | Female | Caucasian | dcSSc | 3.4 | 32 | 2 |
| d | 45 | Female | Caucasian | dcSSc | 5.0 | 20 | 2 |

Age, disease duration, mRSS, and local skin score were recorded at the time of biopsy. Local skin scores were taken adjacent to the site of the biopsy.

