## Supplementary material for "Deletion of adipocyte Sine Oculis Homeobox Homolog 1 prevents lipolysis and attenuates skin fibrosis": Review comments

**Manuscript number:** RC-2024-02521

### 1. General Statements

We sincerely appreciate the time and effort that the reviewers have dedicated to reviewing our manuscript. The insightful comments have highlighted critical areas for improvement, and we are grateful for their constructive feedback. We are pleased that the reviewers recognized the significance of our findings and the potential impact on understanding the role of SIX1 in skin fibrosis and systemic sclerosis (SSc). We have carefully reviewed the comments from the reviewers and we have a developed a plan to enhance the clarity and depth of our research.

In response to reviewers’ comments, we have incorporated many of the recommendations and need for clarification that were identified. We have also included planned experiments that aim to provide a direct link between SIX1 and the downstream mediator PAI-1. We have also discussed technical limitations that would prevent us from performing some of the analysis that were suggested. We would also like to take this opportunity and address some of the comments regarding the novelty of the manuscript. As R1 has pointed out, our group has identified a role for epithelial SIX1 expression in pathophysiology of lung fibrosis through MIF expression. However, herein we demonstrate a unique role for adipocyte SIX1 expression in pathophysiology of dermal fibrosis. These observations differ from our published studies in the lung since they demonstrate a role for SIX1 in a completely different cell: the adipocyte in a completely different disease namely scleroderma where we report on the role of SIX1 on skin fibrosis. Furthermore, the role of adipocyte SIX1 does not appear to involve previously published downstream mediators TGFb and MIF for which we have now included data.

### 2. Description of the planned revisions

R1:

1. Figure 5: The effective depletion of Six1 has been shown only through RT-PCR. Validation with PCR on genomic DNA would be more convincing.
   a. The UBC-Cre line is very hard to breed, and we no longer have the line available however we are in the process of treating our iAdiponectinCre SIX-/- line with tamoxifen and developing a protocol to isolate the DWAT layer to identify SIX1 depletion by gDNA.
2. The claim in Figure 4, "that SIX1 levels are increased in adipocytes by day 7 of bloo" is unconvincing and unsupported, and needs quantification, particularly with data from a mouse model.
   a. We have now included SIX1 RT-qPCR data on Figure 3 (day 7 time point) and we will be performing morphometric assessment of increased SIX1 signals in adiponectin positive cells to further support our findings.

R2:

1- Additional insight into mechanism-for example by clearly uncovering mechanistic insights such as novel downstream targets-is essential.
   a. Our supplementary figure 3 identified potential downstream targets for SIX1 deletion in 3T3-L1 cells treated with a differentiation cocktail into adipocytes. Herein we identified PAI-1 encoded by the gene Serpine1 as a potential target. PAI-1 is indeed elevated in our Bleo-treated mice and depleted in mice lacking SIX-1 in adipocytes. To further identify a link between SIX1 and PAI-1 we plan on performing a luciferase reporter assay linking SIX1 upregulation with SERPINE-1 expression.

R3

1- 4.The last paragraph of the results section about the expression levels of Six1 and Serpine1 in the two patient cohorts is speculative. Both datasets are obtained from whole skin including different cell types. Upregulation of both genes in this context doesn't necessarily mean that they are functionally or mechanistically connected. One gene might be mainly upregulated in one cell type (e.g. adipocytes) the other one in another cell type (e.g. fibroblasts, leukocytes, epidermis). This issue should be either verified by additional experiments (e.g. analysis of co-localization) or the paragraph removed.
   a. We agree with this statement and we will include staining for PAI-1 in SSc skin samples to validate these observations and provide localization information.

### 3. Description of the revisions that have already been incorporated in the transferred manuscript

As mentioned in the general statements, we have made an effort in incorporate the majority of the recommendations identified by the reviewers. We have identified them point-by-point below

**R1:**

1. The abstract section is very long and should be focused on information relevant to the results/discussion. Likewise, the discussion is also too long and could be shortened.
   a. We have revised the abstract and discussion based on these recommendations.
2. Previous work in this group has identified the same gene (SIX1) as a mediator in lung fibrosis, and in the present manuscript, its role is investigated in a subtype of skin cell, without introducing much novelty. Most of the results are poorly investigated; they should be quantified and interpreted with the use of statistics
   a. We have addressed the concerns regarding novelty in section 1. In addition, we have clarified the use of statistics in both the methods section and in each figure legend.
3. The manuscript lacks sufficient quantification and statistical evaluation of the results presented. Specifically, the levels of PAI1, a key SIX1 target, in PRESS and GENISOS cohorts, as well as in cohorts shown in Figures 1 and 2, should be quantified. The quantification of data presented, as well as statistical evaluations, should be applied consistently across all figures.
   a. As mentioned above all figure legends now incorporate clear statistical tests used to analyze the results.
4. In Figure 1, "The bioinformatic analysis identified genes that are expressed at comparatively higher levels in a specific cell type and created a signature score for each cell type being evaluated". Please clarify the bioinformatic analysis used in Figure 1C and specify the tool used for the GO analysis in Figure 1E.
   a. This has been clarified both in the methods and figure legends
5. In Figure 2, the legend mentions " Representative images of dual in situ hybridization for SIX1 (teal) and FABP4 (pink) in 15 SSc and 5 demographically-matched control biopsies". However, Supplementary Table 4 lists only 4 samples (1 control and 3 SSc).
   a. This has been corrected to reflect the correct number of samples used.
6. Figure 3 mainly describes fibrosis in the skin model and is introductory to Figure 4. For a better understanding, the Authors should combine these Figures. The Authors should provide levels of SIX1 (and PAI1) RNA and by IHC in bleomycin-treated skin.
   a. Six1 levels are now included in Figure 3 in addition to SERPINE1. PAI-1 IHC is present in figure 10. Since we are adding morphometric evaluation for Six1 signals in adipocytes, this data has now been added to the revised Figure 4.
7. Figure panel 3c: Please clarify how many mice were used
   a. This has been clarified in the figure legends an N of 5 per time point was utilized
8. Figure 5: SIX1 can be measured by PCR in untreated skin, and therefore, SIX1 mRNA levels should be shown for all the relevant figures. A minor concern is raised for the PCR analysis where the levels of SIX1 calculated from the means in the panel (reported as 2^-∆Ct) are 0.9993 in tamoxifen-KO versus 0.9958 in the WT. What was the number of samples and the CT used for measuring such (statistically relevant) difference? How many times was the PCR repeated?
   a. We have added SIX1 levels for Figure 3. Regarding SIX1 levels in Figure 5b., the N number are as follows: N=9 for iUBCCre and N=7 for iUBC-SIX1-/- groups of which 4 did not express any *Six1*.
9. Figure 5: The "gene expression profiling" analysis in mice in Figure 5 includes only a limited number of genes and should be expanded to include a more comprehensive panel (>4 genes).
   a. We have included additional genes: Acta2, Pparg, adiponectin, Cepba and Mif.
10. The levels of SIX1 observed in WB of 3T3 L1 cells are substantial and not completely silenced, the authors should comment on this.
    a. This has also been discussed by R2 and R3 where complete depletion by siRNA is not usually achieved. Although we include WB quantification for SIX1 siRNA in Supplementary Figure 2, we have also included it for Figure 10.
11. Please define DWAT, AMT and FABP4 in the main text.
    a. Thank you for identified this, these have been defined.

**R2:**

1- I am not sure why the authors focused on serpine, when it appears that TGFb1 and tgfbeta1 cohort is reduced. My suspicion is that the data are due to reduced tgfbeta expression. This can be assessed using rt PCR, histology or ELISA.
   a. We appreciate the reviewer’s insight into the tgfbeta mechanism. We have now incorporated RT-qPCR data for Tgfb1 on day 7 of SQ Bleo (New Figure 3 i), in Bleo treated iUBC-SIX1-/- mice (New Figure 5l) and in our in vitro experiments in 3T3L1 cells with SIX1 KD (Supplementary Figure 2f). Although we report elevated Tgfb1 on day 7 of SQ Bleo that is then inhibited in mice lacking SIX1, our mechanistic studies do not seem to link elevated SIX1 with increased Tgfb1 expression (Supplementary Figure 2f). However, our newly incorporated Serpine1 expression levels (New Figure 3j and Figure 5m) are in line with our mechanistic studies showing increased expression at day 7 of SQ Bleo followed by reduced expression in mice lacking SIX1.
2- Given the effect on fat, it is perhaps surprising that pparg was not examined, especially given the role of pparg in the bleomycin model.
   a. Our mechanistic studies in supplementary figure 2 do not point at SIX1 as regulator of pparg. This is in line with our rt-pcr data for Pparg, showing no changes on day 7 of bleo treatment or following deletion of Six1 using the UBC promoter. These results indicate that SIX1 does not alter Pparg expression.

**R3**

2- 1.In the text, the authors state that SIX1 expression levels were elevated in both lcSSc and dcSSc. In the corresponding Fig. 1b, however, the increase in lcSSc looks very small (which might not have biological relevance) and no significance level is shown. Could the authors clarify this issue?
   a. The significance level is P<0.001 and this is based on a N number of 43lcSSc vs 44 controls. This represents a robust significance and N number. These have been clarified in the Figure, the Figure legend and the methods. The biological significance is demonstrated by subsequent experiments in this manuscript, primarily in our SIX1 KO studies in mice.
3- 2.The authors write in the text about the ubiquitous knockout of Six1 in mice that they did not observe a difference in dermal thickening upon bleomycin challenge between knockout and "wildtype" mice. However, adipocyte-specific knockout is reducing dermal thickness as shown in Fig. 7. How do the authors explain this phenomenon? A paragraph to the discussion should be added.
   a. Indeed, this is a valid observation that has been addressed in the discussion section. It is either due to more effective deletion of SIX1 using the adiponectin Cre or as a result of a protective effect of SIX1 in another cellular system.

**R3 Minor Comments:**
1. According to methods and the scheme in Fig. 3a, the authors used two different sites on the back (upper and lower back) of the mice for bleomycin injections. Two questions are arising that should the authors clarify: Did the authors analyse both sites seperately or did they pool them (similarly to humans, skin architecture and thickness differs between sites)? How did the authors avoid skin reactions (e.g. scar formation) due to repeated injections at the same sites?

The upper and lower back samples were not pooled, instead we used the upper sample for RT-qPCR and the lower sample for histology. We have clarified this in the methods. As for scar formation, our PBS mouse studies that represent the control for Bleo injections do not reveal significant changes in fibrotic injury.

2. Ref. 42 is missing in the reference list. Please add.

We have corrected this

4. In Fig. 9, in my opinion, the pictures shown do not reflect the strong difference shown in the quantification graph. Maybe that could be adjusted.

- We have enhanced the contrast of the figures to highlight the differences more clearly

5. R3 Minor Comment 3. Microscopic pictures showing cell morphology would add to the information given in Supp. Fig.2.

a. We agree that this could provide further information regarding the differentiation state of the cells, we now include Oil O red staining showing that the cells were indeed differentiated into adipocytes (Supp Fgure 2g)

### 4. Description of analyses that authors prefer not to carry out

5. R1: In Figure 2A; The amount of DWAT is evaluated only using H&E staining. Performing Red Oil staining would be more convincing.
   a. We agree that this method would highlight adipocytes, however, since we have collected FFPE samples, oil-o-red is not compatible with this fixation method. In addition, our studies are based on previous publications that used masson’s trichome (what we used) to identify the DWAT layer in mouse tissues
      i. Arthritis Rheumatol 2015 Apr;67(4):1062-73. doi: 10.1002/art.38990.
6. R1: Why are IHCs not used with antibodies instead of in situ hybridizations? The Protein Atlas repository for single-cell analysis shows expression of SIX1 in fibroblasts, myoblasts, and endothelial cells (adipose cells are not present, but it might be a consequence of the markers used to identify the subpopulations). The absence of SIX1 expression in other cell subtypes in in situ analyses, even after fibrotic insult, needs clarification.
   a. We have selected RNAscope over IHC for the following reasons:
      i. RNAscope is more selective than IHC were performance if often LOT number based and there is cross-reactivity between proteins such as for SIX1 and SIX4.
      ii. As identified by the Human Protein atlas, SIX1 is an important transcription factor in development that is usually expressed in cancer and disease processes. Thus, it is possible for SIX1 to be expressed in skin endothelial, fibroblasts and myofibroblasts during developmental processes that are outside the scope of this manuscript. The observation that SIX1 is elevated primarily in adipocytes in our hands provides further rationale to the study its role in this cell type in the context of dermal fibrosis.
7. R2 siRNA knockdown is a reasonable approach, as long as a "smartpool" or more than one siRNA is used AND knockdown verified at a protein level.
   a. We agree that the “smartpool” approach is now commonly performed. Our selection of the siRNA is based on previous studies (JCI Insight 2022 May 23;7(10):e142984) where we selected an efficient siRNA approach. We do not think that the addition of a smartpool approach to contribute further to the present study
8. R3: With Six1 with a quite specific expression in adipocytes and very low expression in fibroblasts, it is confusing that the authors knocked down Six1 in 3T3 fibroblasts in Supp. Figs. 2-3 and Fig. 10. In my opinion, it would make more sense to knock down Six1 in adipocytes and analyse potential adipocyte-myofibroblast-transition [AMT].
   a. We agree with R3 in that SIX1 deletion in adipocytes to evaluate AMT would be ideal. However, culturing adipocytes directly is not possible since they must be differentiated first from 3T3-L1 cells first. Next it is very challenging to perform KD studies in adipocytes as such our studies were designed to evaluate the capacity of SIX1 to regulate cell identity since 3T3-L1 cells were exposed to a differentiation cocktail to promote adipocyte differentiation. Surprisingly, these studies demonstrate that SIX1 deletion in 3T3-L1 cells did not affect adipocyte markers (Adiponectin, Cebpa or Pparg) but were able to alter expression of profibrotic mediators including PAI-1 encoded by Serpine1. These findings suggest that SIX1 deletion does not alter fibroblast to adipocyte transition and thus it is unlikely that it mediates AMT. However, our results point at the capacity of SIX1 in promoting the expression of profibrotic mediators, namely PAI-1.
9. R3: 5. Optional: The authors show increased levels of Six1 upon bleomycin challenge in mice, but we do not learn what is the initiating trigger for this (also for human samples). As the bleomycin model is an inflammation-driven model, infiltrating leukocytes might be part of the process. The authors could more investigate that and/or add a paragraph to the discussion.
   a. We agree with R3 in that leukocytes infiltration could drive the expression of SIX1 in SSc and Bleo treated mice, however uncovering this process is likely to require experiments outside the scope of this manuscript.
10. R3: 6. Optional: For future study of mechanisms and translational impact, it might be helpful to include another inflammation-independent animal model of fibrosis (e.g. overexpression of TBRIact).
    a. We agree that an additional mouse model would enhance our study, yet many of these models are difficult to obtain and would require many months to complete placing them outside the scope of our research.
